# Genetic Basis of *De Novo* Appearance of Carotenoid Ornamentation in Bare-Parts of Canaries

**DOI:** 10.1101/762112

**Authors:** Małgorzata Anna Gazda, Matthew B. Toomey, Pedro M. Araújo, Ricardo J. Lopes, Sandra Afonso, Connie A. Myers, Kyla Serres, Philip D. Kiser, Geoffrey E. Hill, Joseph C. Corbo, Miguel Carneiro

**Author notes:** contributed equally (Małgorzata Anna Gazda, Matthew B. Toomey). Corresponding authors: Małgorzata Anna Gazda, Matthew Toomey, Miguel Carneiro.

## Abstract

Unlike wild and domestic canaries (*Serinus canaria*), or any of the three dozen species of finches in genus *Serinus*, the domestic urucum breed of canaries exhibits bright red bills and legs. This novel bare-part coloration offers a unique opportunity to understand how leg and bill coloration evolve in birds. To identify the causative locus, we resequenced the genome of urucum canaries and performed a range of analyses to search for genotype-to-phenotype associations across the genome. We identified a nonsynonymous mutation in the gene *BCO2* (beta-carotene oxygenase 2, also known as *BCDO2*), an enzyme involved in the cleavage and breakdown of full-length carotenoids into short apocarotenoids. Protein structural models and *in vitro* functional assays indicate that the urucum mutation abrogates the carotenoid cleavage activity of BCO2. Consistent with the predicted loss of carotenoid cleavage activity, urucum canaries had increased levels of full-length carotenoid pigments in bill tissue and a significant reduction in levels of carotenoid cleavage products (apocarotenoids) in retinal tissue compared to other breeds of canaries. We hypothesize that carotenoid-based bare-part coloration might be readily gained, modified, or lost through simple switches in the enzymatic activity or regulation of *BCO2* and this gene may be an important mediator in the evolution of bare-part coloration among bird species.

## INTRODUCTION

Since Darwin and Wallace, the hue and pattern of bird coloration has been a central arena for testing theories of signaling, assessment, and speciation (1–3). The pigmentary and structural bases of avian coloration are now well understood (4). The key missing piece to the puzzle of how and why conspicuous coloration evolves in birds is a better understanding of the genetic and molecular mechanisms controlling the production of color (5, 6). Recent breakthroughs have begun to reveal the genes that control carotenoid pigmentation, which mediates most of the conspicuous red, orange, and yellow coloration in birds. *CYP2J19*, a cytochrome p450 gene, is implicated in the conversion of yellow to red carotenoids and consequently for much of the red coloration in birds (7, 8). *SCARB1* plays a key role in carotenoid uptake and consequently in the presence or absence of coloration (9). Here we add to a growing understanding of the genetic basis of coloration in birds by revealing the mechanism for the *de novo* evolution of red bill and leg coloration in canaries.

Domesticated canaries are derived from the island canary (*Serinus canaria*), a small finch with dull yellow carotenoid-based feather coloration and no carotenoid coloration of bills or legs (10). Several centuries of selective breeding produced domestic canary birds with a fantastic diversity of yellow and red feather coloration, but until very recently no breed of canary had been produced that deposited carotenoids in bills or legs (11). Indeed, none of the three dozen species of finches in the genus *Serinus* have red or yellow leg or bill coloration (12). Several decades ago, a canary with a red bill and legs appeared spontaneously in a colony of red factor canaries, a breed with bright red plumage coloration. Over subsequent years, this mutant canary founded a new breed of canary with bright red bills and legs, now called urucum canary (Fig 1). The pigmentation in bare parts of urucum canaries follows an autosomal recessive pattern of inheritance. Thus, urucum canaries present a unique opportunity to identify a single genetic locus that enables birds to express a conspicuous new ornamental color trait.

**Figure 1.**
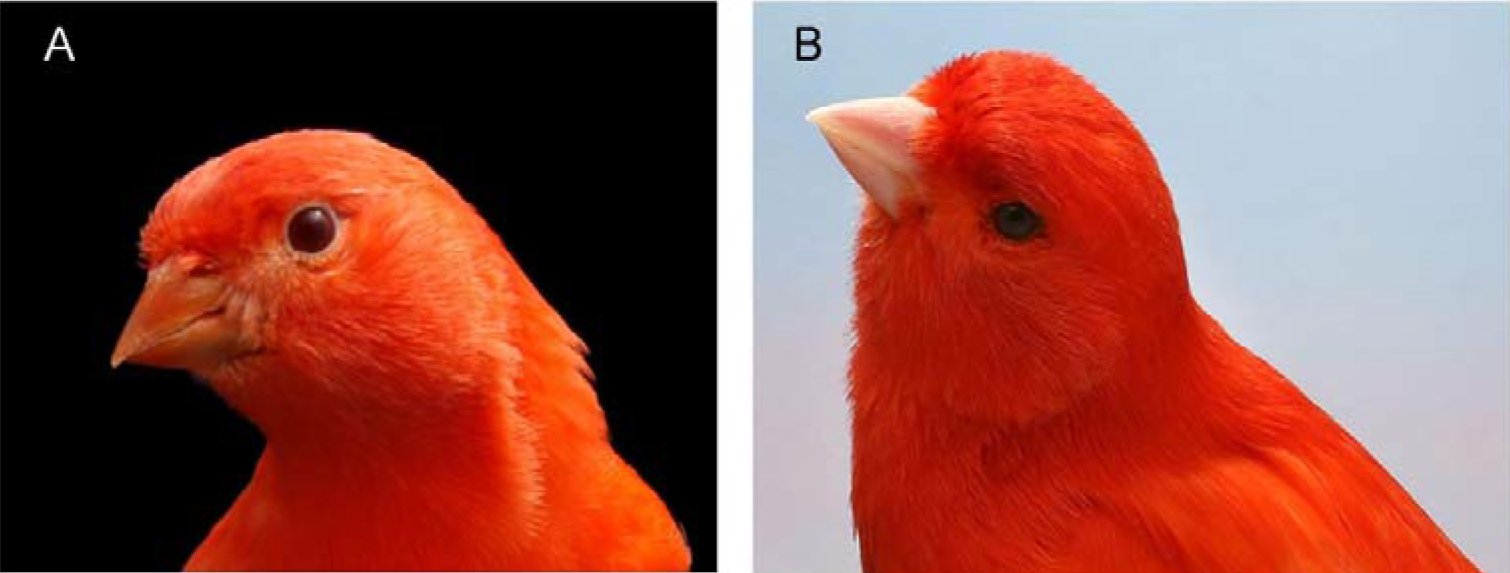
Representative picture of **(A)** urucum canary and **(B)** lipochrome red canary.

To identify the causative locus responsible for the expression of carotenoid-based pigmentation in bare parts in canaries, we resequenced the genome of urucum individuals and performed a range of analyses to search for genotype-to-phenotype associations across the genome. We identified a nonsynonymous point mutation in the gene *BCO2*, an enzyme that catalyzes the oxidative cleavage of both pro-vitamin A and non-pro-vitamin A carotenoids (13) in birds and other vertebrates. We found a perfect association between presence/absence of the *BCO2* mutation and bare-part coloration in canaries. Through protein structural models and *in vitro* functional assays we confirm causality of the mutation and its effect on protein function.

## RESULTS

### Carotenoid content of urucum canary tissues

The feathers and bare parts of the urucum canaries are brilliantly colored. To determine the pigmentary basis of the urucum phenotype, we used high-performance liquid chromatography (HPLC) to examine and compare carotenoid accumulation in the beak and feather tissue of urucum and wild-type canaries with the same plumage color backgrounds. Canaries carrying the urucum mutation tended to accumulate significantly higher concentrations of carotenoids in their beak tissues than wild-type birds (Mann–Whitney U = 0, n_1_ = 6, n_2_ = 3, p = 0.024, Figs 2A, S1-2, Table S1). The carotenoid concentration in the feathers of red canaries was significantly higher than in the feathers of yellow birds (Mann–Whitney U = 0, n_1_ = 5, n_2_ = 4, p = 0.016, Figs 2B, S2-3, Table S2). Feather carotenoid concentration did not differ significantly between urucum and wild-type birds (Mann–Whitney U = 5, n_1_ = 6, n_2_ = 3, p = 0.38, Fig 2B, Table S2).

**Figure 2.**
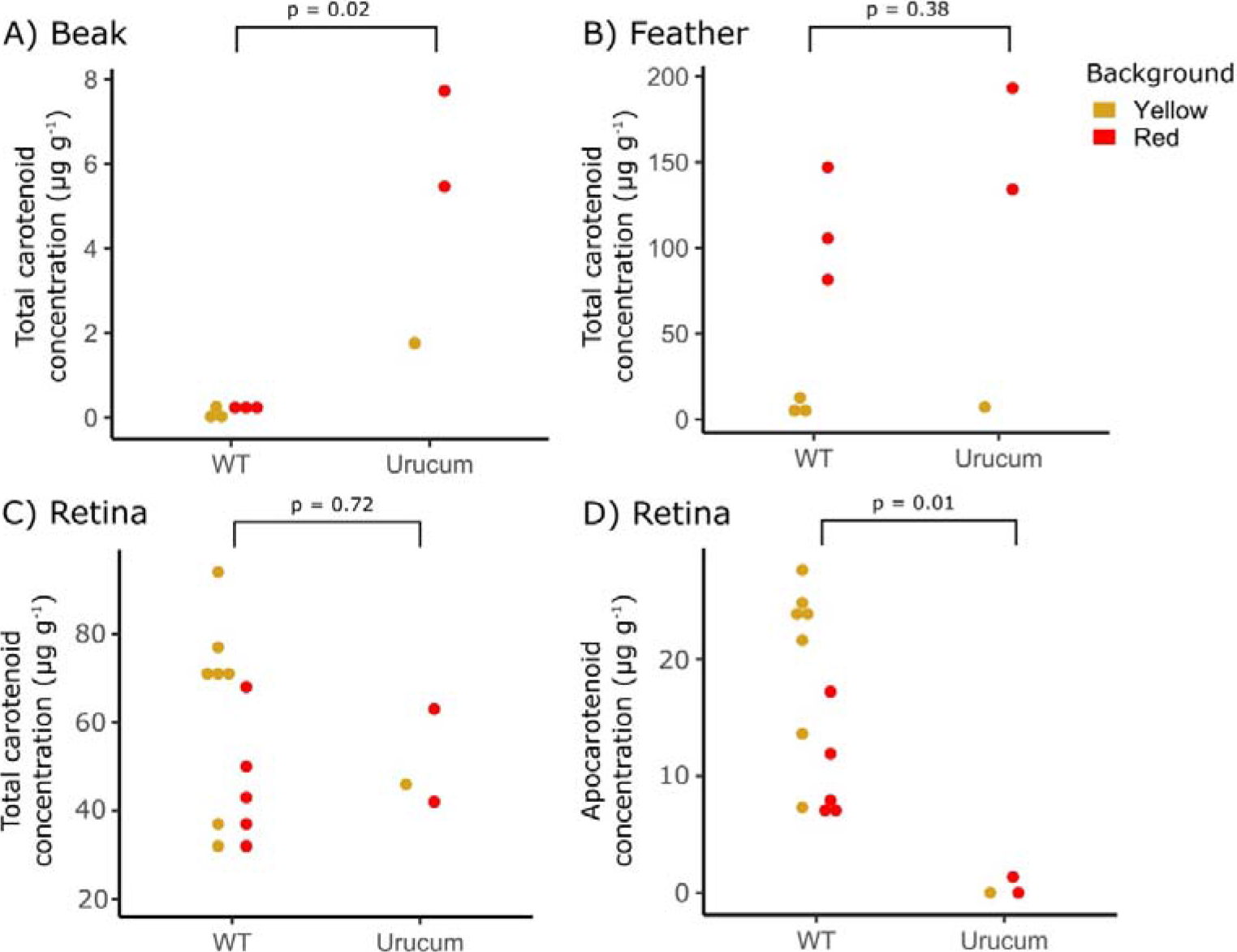
Tissue carotenoid accumulation patterns among urucum and wild-type canaries. Points indicate the value for each individual sampled. The concentration of total carotenoids per gram of tissue are shown for **(A)** beak and **(B)** feathers of wild-type (WT) and urucum canaries with red-factor (denoted in red) or yellow lipochrome (denoted in yellow) genetic backgrounds. **(C)** The concentration of total carotenoids in the retina per gram of total protein sampled. **(D)** The concentration of carotenoid cleavage products, apocarotenoids, in the retina per gram of total protein sampled. Details of the specific carotenoid composition of each tissue are available in Tables S1-3 and Figs S1-5.

A diverse suite of carotenoid pigments accumulate within the oil droplets of the cone photoreceptors of the avian retina where they modify spectral sensitivity and fine tune color discrimination (14–16). To determine if and how the urucum mutation influences carotenoid accumulation in the retina, we used HPLC to examine and compare the retinal carotenoid profiles of urucum and wild-type canaries. The total accumulation of all carotenoid types in the retina did not differ significantly between urucum and wild-type birds (Mann–Whitney U = 21, n_1_ = 12, n_2_ = 3, p = 0.72, Fig 2C, S4-5, Table S3). However, when we examined specific components of the retinal carotenoid profile we found that urucum canaries had significantly lower concentrations of apocarotenoids in their retinas than wild-type birds (Mann–Whitney U = 0, n_1_ = 12, n_2_ = 3, p = 0.011, Fig 2D). Apocarotenoids (e.g. galloxanthin and dihydrogalloxanthin) pigment the cone oil droplets of the blue (SWS2) cone photoreceptor and the principle member of the double cone, and are hypothesized to be produced through the BCO2-mediated cleavage of dietary zeaxanthin (17). This result suggests that urucum birds may have decreased carotenoid cleavage activity. Both wild-type and urucum red-factor background canaries accumulated canthaxanthin in their retinas (Fig S6), which is not typically a component of the avian retinal carotenoid profile (14, 15, M. Toomey unpublished data). In some urucum individuals (fed normal diet with supplement as regular red birds), canthaxanthin constituted as much as 80% of the carotenoid content of the retina (Table S3, Figs S4-6).

### High-resolution mapping of the urucum phenotype

To investigate the genetic basis of bare-part coloration in urucum canaries, we performed whole genome sequencing of a DNA pool of individuals of this breed (n = 20) to an effective coverage of 26X (Table S4). Since the urucum breed was created through the fixation of a spontaneous mutation that emerged in red factor canaries (18), we further used whole-genome resequencing data of a DNA pool of red lipochrome birds (n=16, coverage = 17X) from a previous study (7).

The urucum phenotype is transmitted in a manner consistent with a single recessive allele. Therefore, the genomes of both breeds are presumably very similar except in a single region controlling bare-part coloration. To search for regions of high differentiation between both breeds, we summarized allele frequency differentiation across the genome (scaffolds > 20 kb) using the fixation index (*F*_*ST*_) and a sliding window approach (Fig 3A). We found that levels of genetic differentiation were low to moderate throughout most of the genome (mean *F*_*ST*_ ~ 0.15), but that one region displayed very high levels of *F*_*ST*_ when compared to the remainder of the genome. This region contained the top 36 values of the empirical distribution of *F*_*ST*_ (range= 0.62-0.89) and spanned a region of ~200 Kb (715,000 to 930,000 bp) on scaffold NW_007931177, which is homologous to zebra finch chromosome 24. Using a backcross mapping population (see Methods), we found a perfect association between our candidate region and bare-part coloration: all wild-type individuals (n=11) were heterozygous for a SNP diagnostic between the urucum and wild-type parental individuals, and all birds exhibiting the urucum phenotype (n=7) were homozygous for the allele present in the parental urucum individuals.

**Figure 3.**
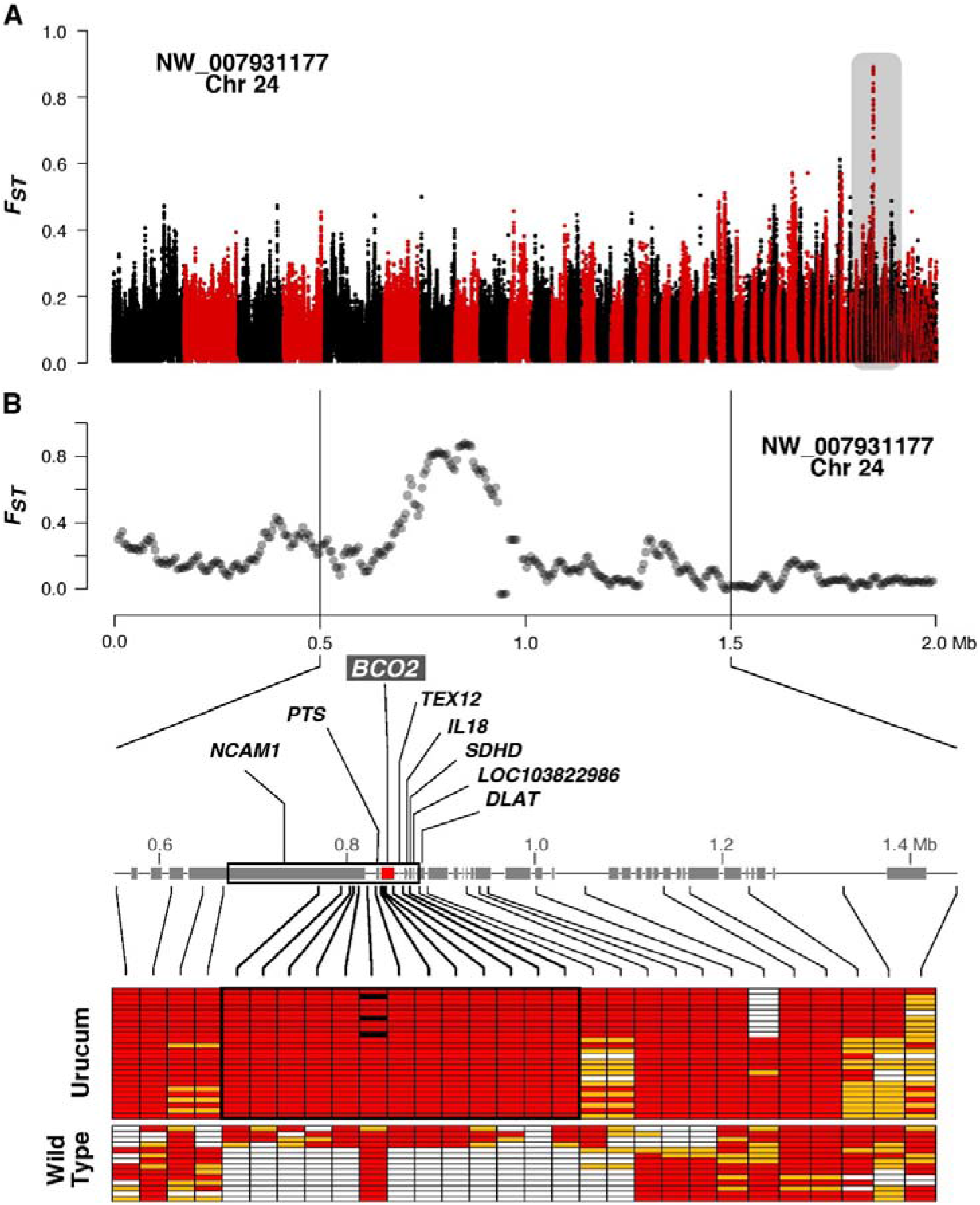
Mapping of the urucum mutation. (A) Selective sweep mapping. *F*_*ST*_ between urucum and lipochrome red breed across the genome. Each dot represents *F*_*ST*_ in 20-kb windows iterated every 5 kb. The different scaffolds are presented along the *x* axis in the same order as they appear in the canary reference genome assembly. **(B)** *F*_*ST*_ zoom-in and IBD mapping. (Top) *F*_*ST*_ in 20-kb windows iterated every 5 kb across the outlier region (delineated by vertical lines). (Bottom) The protein-coding genes found within this region are indicated by grey boxes. For the IBD analysis, 30 SNPs were genotyped for 27 urucum canaries and 14 individuals belonging to three breeds with yellow, red and white coloration. Alleles more common in urucum canaries are represented by red boxes, alternative alleles are represented in white, and heterozygous are represented in orange. The black-outlined box indicates a segment of high homozygosity in urucum canaries and black boxes indicate missing data.

We next carried out identity-by-descent (IBD) mapping (Fig 3B) to increase the resolution at the candidate locus, under the assumption that the urucum mutation emerged a single time within a haplotype that should be shared by all urucum individuals. We genotyped 30 variants selected from the whole-genome resequencing data in a larger cohort of birds, including urucum individuals (n = 27) as well as individuals (n = 14) belonging to several breeds lacking coloration in bare parts. Unlike for the other breeds, we found that urucum canaries were homozygous for a large continuous genomic segment defined by 13 SNPs, spanning a physical interval of 104 kb (770,364-874,443) that harbored eight protein-coding genes: *NCAM1*, *PTS*, *BCO2*, *TEX12*, *IL18*, *SDHD*, *LOC103822986*, *DLAT* (Fig 3B; Table S5). Of these, the *BCO2* represents a strong candidate for the gene underlying the urucum phenotype since it encodes an enzyme that is known to cleave carotenoids (19, 20) and has been previously implicated in carotenoid pigmentation of the integument in birds and reptiles (21–24).

### An amino acid substitution at a universally conserved position in *BCO2* is perfectly associated with the urucum phenotype

We next screened the IBD interval for potential causative mutations using the whole-genome resequencing data. One single point mutation clearly stood out at nucleotide position 837,806 that was predicted to be a nonsynonymous mutation in the exon 9 of the *BCO2* gene. This variant results in the substitution of a histidine for an arginine at residue 413 of the protein (R413H) that could potentially affect *BCO2* activity (see below). We genotyped the nonsynonymous variant in the cohort of urucum samples used for the IBD analysis above, and consistent with an autosomal recessive mode of inheritance, we found that all 27 individuals were homozygous for the allele encoding the amino acid arginine. Interestingly, domesticated canaries expressing yellow carotenoids in their bare parts (n=4), which have emerged more recently, were also homozygous for the same nonsynonymous variant, suggesting the same genetic basis for bare-part coloration in both red and yellow birds. A multispecies alignment of the *BCO2* protein, including 175 species from all major vertebrate groups revealed that this amino acid is universally conserved (Fig 4A, Fig S7).

**Figure 4.**
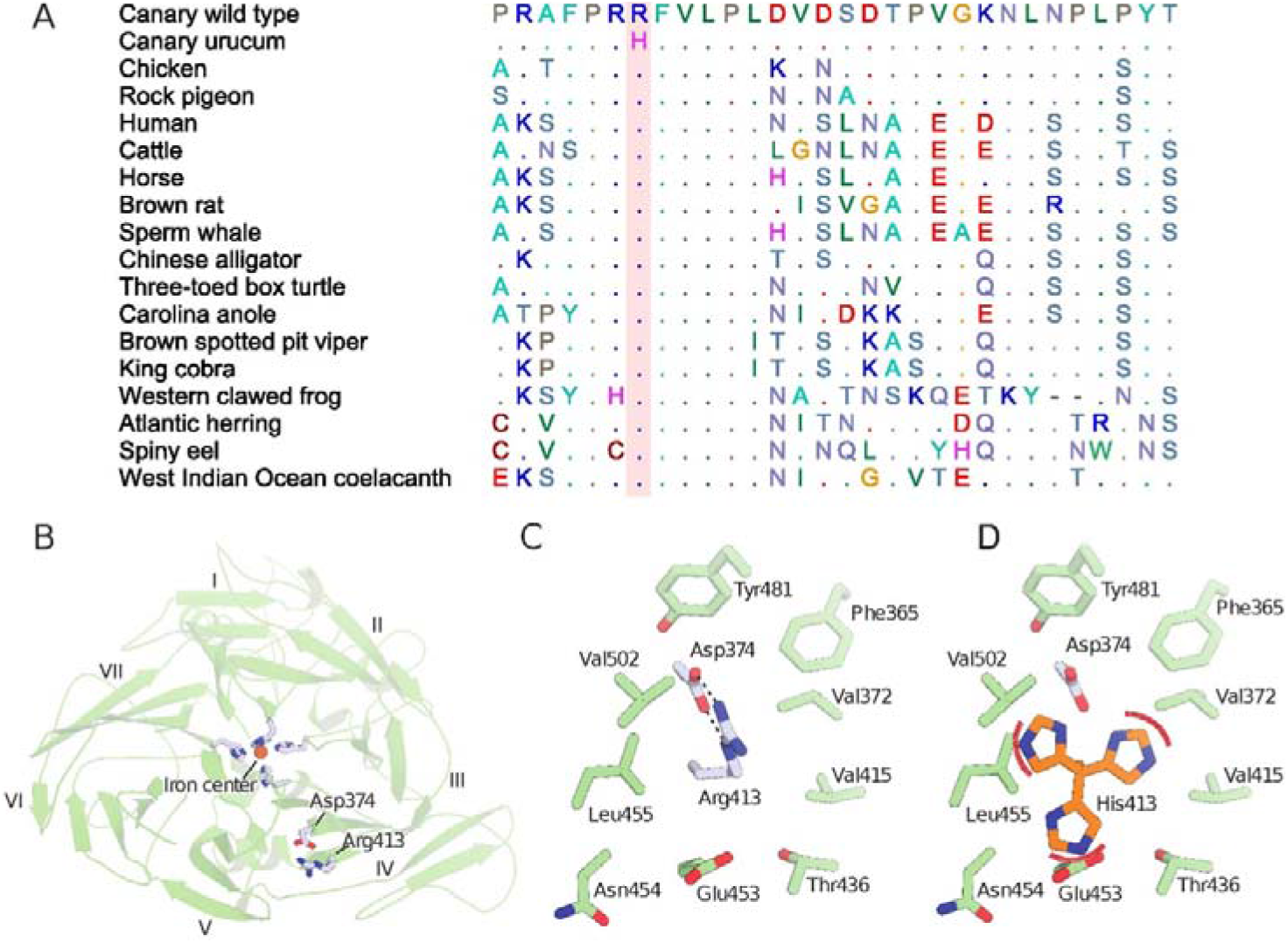
Structural consequences of the R413H substitution in canary *BCO2*. **(A)** Multi-species alignment of vertebrates surrounding the candidate missense mutation in the highly conserved exon 9 of *BCO2* (the missense mutation is highlighted in red).The dots represent the same nucleotide as the reference wild-type canary allele at a given position. **(B)** Homology model of canary *BCO2* based on an RPE65 template. Blades are numbered according to convention. **(C)** Close-up view of R413 forming a twin salt bridge interaction with Asp374 within a pocket composed of predominantly non-polar side chains. **(D)** Substitution of Arg413 with His results in predicted steric clashes with surrounding side chains that would disrupt the protein structure or cause a protein folding defect.

To help define the structural consequences of the R413H substitution in *BCO2*, we first generated a homology model of canary *BCO2* using a structurally related protein, RPE65, as a template (Fig 4B). Inspection of the model revealed that Arg413 is predicted to form a twin salt bridge with Asp374 at a location distant from the iron center of the enzyme. The same type of interaction is formed between equivalent residues of other members of the carotenoid cleavage oxygenase family of enzymes, including *Synechocystis* ACO (25) and bovine RPE65 (26), indicating that it is evolutionarily conserved and structurally or functionally important. Indeed, Arg-carboxylate salt bridges are known to contribute substantially to protein thermal stability (27). The Arg-Asp pair is in a hydrophobic pocket within blade IV of the beta-propeller domain and the interaction appears important for maintaining an overall neutral charge in this apolar environment (Fig 4C). *In silico* substitution of His for Arg at position 413 resulted in severe steric clashes in all rotameric orientations of the His residue that could not be resolved by secondary rotamer changes in the involved residues (Fig 4D). This observation suggests that the R413H could cause structural distortion of the protein, leading to a possible loss of activity. Alternatively, the substitution could cause a gross protein-folding defect via these steric clashes or through electrostatic destabilization. Overall, the striking conservation of the amino acid position, together with the structural modelling of the protein sequence, strongly suggests that this mutation is functionally relevant.

Finally, we evaluated whether regulatory mutations could also be implicated in the urucum phenotype. One way to test this possibility is to study allele-specific expression in the shared cellular environment of individuals heterozygous for the urucum and wild-type alleles. If *cis*-acting regulatory mutations are driving expression differences, then one allele should be expressed preferentially. We sampled one heterozygous individual and carried out cDNA sequencing. Using the nonsynonymous position as a marker for allelic identity, we found that the number of reads containing the wild-type and urucum alleles was similar (48% *vs.* 52%, out of a total 193,155 reads overlapping the exon of this gene). We therefore conclude that there are unlikely to be any functionally significant *cis*-regulatory differences between the two alleles.

### Functional analysis shows that the *BCO2* mutation results in compromised enzymatic activity

To determine if and how the candidate causal variant altered the carotenoid-cleavage activity of BCO2, we evaluated the ability of wild-type and urucum BCO2 to cleave zeaxanthin and canthaxanthin substrates. We found that bacterial lysates containing recombinantly expressed wild-type BCO2 readily cleaved both carotenoid substrates and generated apocarotenoid products (Figs 5 and S8). In contrast, lysates containing the urucum BCO2 variant, with the R413H substitution, did not produce any detectable apocarotenoids (Figs 5 and S8). Additionally, the reaction containing the mutant BCO2 variant retained substantially more substrate than the wild-type reaction (Fig 5). In reactions with the zeaxanthin substrate, small amounts of apocarotenoid were detectable in all conditions, including assays with heat-denatured lysates (Fig S8). We therefore believe that these trace amounts of apocarotenoid may be a product of non-enzymatic oxidation (28). Our analysis indicates that the urucum mutation eliminates the enzymatic activity of BCO2.

**Figure 5.**
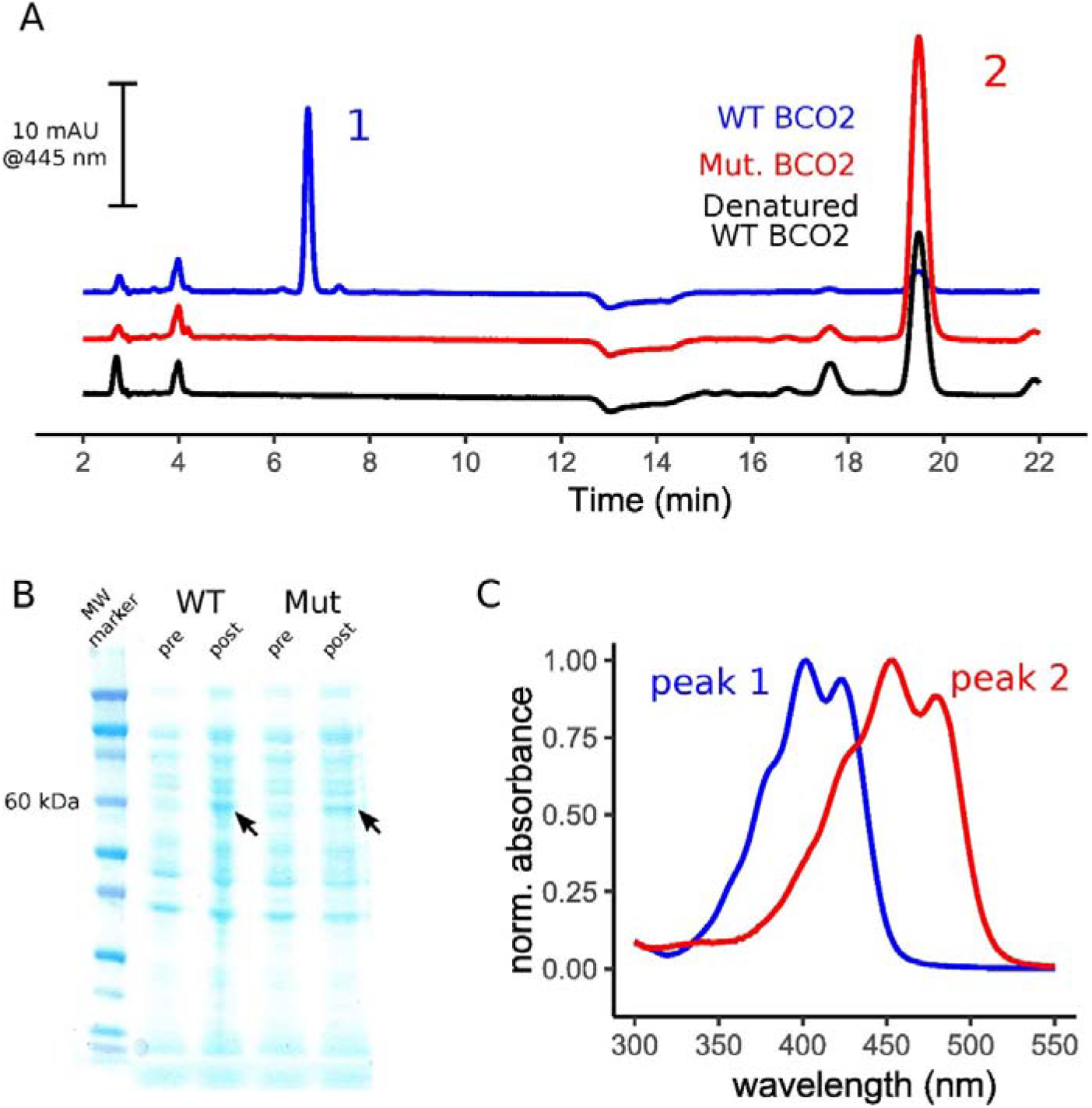
*In vitro assay* of the cleavage activity of urucum and wild-type variants of *BCO2*. **(A)** A Coomassie-stained SDS-PAGE gel visualizing protein expression in the BL21 DE3 cultures before (pre) and after (post) the induction of recombinant protein expression. Arrowheads indicate the enrichment of an ~60 kDa band that is consistent with the induction of the expression of the recombinant wild-type (WT) and mutant (Mut) *BCO2*. **(B)** Representative HPLC chromatograms of carotenoids extracted from reactions containing canthaxanthin substrates and wild-type, mutant, or wild-type heat denatured bacterial lysates. **(C)** UV-Vis light absorbance spectra of peaks 1 and 2. Note that these samples have been reduced with sodium borohydride and the canthaxanthin substrate (peak 2) has been reduced to isozeaxanthin (β,β-carotene-4,4’-diol). The short retention time and short-wavelength shifted absorbance spectrum of peak 1 is consistent with an apocarotenoid which we hypothesize to be 10’-Apo-β-carotene-4,10’-ol. We found no evidence of this apocarotenoid in the mutant or denatured *BCO2* assays.

## DISCUSSION

Through genomic and biochemical analyses of the urucum canary breed, we identified a single-nucleotide missense mutation affecting an evolutionarily invariant amino acid that results in *de novo* carotenoid-based coloration of the bill and legs. This mutation in the *BCO2* gene disables the enzyme’s carotenoid cleavage activity, thereby causing a marked increase in the concentrations of intact carotenoids in tissues. The increase in carotenoid pigmentation of beak, legs, and skin of the urucum breed suggests that degradation of carotenoids by *BCO2* is a key mechanism for regulating the presence or absence carotenoid coloration in these tissues in birds and could be an important target of selection in the evolution of color patterns.

Previous studies have implicated BCO2 in species-typical carotenoid coloration. Many breeds of domestic chickens have yellow legs but the wild progenitor of chickens, the red junglefowl (*Gallus gallus*), has gray legs lacking carotenoid coloration. Using QTL analysis (21) mapped the yellow leg coloration phenotype to a genomic region that contains *BCO2*. The authors found a tissue-specific reduction in *BCO2* in the legs of yellow birds, suggesting a cis-regulatory mutation affecting *BCO2* expression, but the causal mutation was not identified. The authors suggested that this mutation had been acquired through introgression from a sister species, the gray junglefowl (*Gallus sonneratii*), which has yellow legs. Subsequently, *BCO2* has been implicated in carotenoid coloration in various animal groups, such as warblers (23) and lizards (24), as well as in control of fat and milk color in domestic animals (29–31). Here we show that a single missense mutation in *BCO2* can lead to large changes in the ornamental color pattern of a songbird. The carotenoid-cleaving activity of BCO2 presents a mechanism by which carotenoid-based bare-part coloration might be readily gained, modified, or lost through simple regulatory switches controlling tissue-specific expression of *BCO2* or changes in its enzymatic activity. This relatively simple mechanism is consistent with the apparent evolutionarily rapid loss and gain of bare-part coloration in many groups of birds (32) and raises the possibility that these colorful traits might be particularly labile through the course of evolution.

In addition to bare-part pigmentation, the retinal carotenoid profiles of urucum canaries differ from wild-type birds, indicating that BCO2 plays an important role in the pigmentation of the cone oil droplets that fine tune the spectral sensitivity of photoreceptors. In the urucum canaries, we observed little or no apocarotenoid pigment in the retina. Apocarotenoid pigments selectively accumulate in the oil droplets of the blue cone (SWS2) photoreceptor and the principal member of the double cone photoreceptor (15, 17). We previously speculated that the selective accumulation of apocarotenoids is mediated by the cone subtype-specific expression of the BCO2 within the blue and double cones (17). The reduction or absence of apocarotenoid pigmentation in the retinas of urucum canaries supports this hypothesis.

Detailed understanding of carotenoid metabolism and its biochemical and physiological importance hold the potential for better understanding of not only ornamental coloration in birds but also the role of micronutrients in health and disease in vertebrates more generally. The pathophysiology of the urucum mutation has not been fully characterized, but breeders have reported that urucum birds suffer from problems with balance, motor abilities, coordination, inactivity and vision. Previous studies have shown that high levels of carotenoid accumulation can have negative physiological effects. For example, in American goldfinches (*Carduelis tristis*), long-term exposure of tissues to high levels of carotenoids resulted in a gradual deterioration of skeletal muscle tissue and consequent defects in flight performance (33). In mouse, *BCO2* knock out results in the accumulation of abnormally high levels of xanthophyll carotenoids in the liver and other tissues (20). *BCO2* mutant mice show evidence of mitochondrial dysfunction, elevated oxidative stress, and liver steatosis, indicating that *BCO2* plays an important role in maintaining carotenoid homeostasis and protecting against the potentially damaging effects of these pigments (20). Similar detrimental effects of *BCO2* knockdown have also been demonstrated in developing zebra fish (*Danio rerio*) and human cell lines, indicating that the role of BCO2 in carotenoid homeostasis is broadly conserved (34, 35). Our observations suggest that the interplay of the signaling benefits of carotenoid coloration and potential physiological costs of concentrations of these pigments in tissues may be mediated by regulation of BCO2.

## MATERIALS AND METHODS

### HPLC analyses in feathers, beak, and retina

We used high-performance liquid chromatography (HPLC) to determine the composition and concentrations of carotenoid pigments in the tissues of two urucum canaries with a red-factor background, one urucum canary with a yellow plumage background, three to five red-factor canaries that were wild-type for the urucum mutation and three to seven yellow lipochrome canaries that were wild-type for the urucum mutation. To assay the carotenoid content of the feathers, we plucked 10 from the nape of each bird and measured the total mass of the feathers sampled from each individual on a laboratory balance. We extracted pigments by incubating the feather for six hours at 65°C in 2 ml of acidified pyridine (36). We then added 2 ml of distilled water, and extracted with 2 ml of hexane:tert-methyl butyl ether (1:1 vol:vol). We collected the hexane:tert-methyl butyl ether fraction, evaporated to dryness under a stream of nitrogen and resuspended in 200 μl of methanol:acetonitrile 1:1 (vol:vol). We injected 50 μl of each extract into an Agilent 1100 series HPLC with a YMC carotenoid 5.0 μm column (4.6 mm × 250 mm, YMC). We separated pigments with a gradient mobile phase consisting of acetonitrile:methanol:dichloromethane (44:44:12) (vol:vol:vol) for 0-11 minutes then a ramp up to acetonitrile:methanol:dichloromethane (35:35:30) from 11-21 minutes followed by a return to isocratic conditions through 35 minutes. The mobile phase was pumped at a constant rate of 1.2 ml min-1 and the column maintained at 30°C for the entirety of the run. We monitored the samples with a photodiode array detector at 400, 445, and 480 nm, and identified and quantified carotenoids as described in Toomey et al. (9).

We measured the retinal carotenoid content following previously published methods (9). Briefly, we homogenized the retinas in phosphate buffered saline and extracted with hexane:tert-methyl butyl ether (1:1 vol:vol). We divided each sample in two and saponified with weak (0.02 M NaOH in methanol) or strong (0.2 M NaOH) base to optimize the recovery of ketocarotenoids or other carotenoid classes respectively. We separated and quantified carotenoids on the HPLC system described above. Chromatographic conditions identical to feather analyses were used for the ketocarotenoid portion of the sample. The strong base samples were run under modified conditions to optimize the separation of apocarotenoids. For these samples, we cooled the column to 18°C and pumped mobile phase at a constant rate of 1.0 ml/min. We used a gradient mobile phase beginning with acetonitrile:methanol (50:50) for 9 min, followed by a ramp up to acetonitrile:methanol:dichloromethane (44:44:12) (vol:vol:vol) from 9–11 min, isocratic through 21 min, then a second ramp up to acetonitrile:methanol:dichloromethane (35:35:30) from 21–26 min followed by isocratic conditions through 35 min.

The extractions of carotenoids from the beak tissue proved challenging and we used two different methods depending upon the color background of the individual. For all individuals, we cut a ~0.5 cm^2^ portion of the lower mandible and measured total wet mass on a laboratory balance. For the individuals with a yellow plumage background, we extracted carotenoids with acidified pyridine as described above for the feather analyses. We attempted pyridine extraction for the individuals with the red plumage background, but found that the yield of carotenoids was poor and the tissue retained its color. As an alternative, we ground the beak tissue for 30 minutes at 4,000 rpm with a bead mill (BeadBug, Benchmark Sci. Edison, NJ) using 10-2 mm zirconia beads in 1 ml of methanol. We centrifuged the resulting homogenate, collected the methanol supernatant, and dried under a stream of nitrogen. We repeated this extraction three times for each sample and pooled the resulting supernatants for each sample. The carotenoid pigments recovered with the methanol extraction were esterified. Therefore, we saponified the samples with weak (red background) or strong (yellow background) base as described for the retina samples. We separated, identified, and quantified carotenoids with HPLC as described for the feather samples. The methanol extraction yielded a mean of 7.3X more carotenoid than the pyridine extraction of the red background samples. In contrast, the methanol extraction of the yellow background samples yielded only 33% of the carotenoid recovered with the pyridine extraction. Therefore, we report the results for the pyridine extraction for yellow background individuals and methanol extraction for the red background individuals. For all tissues we compared carotenoid concentrations with a Welch’s two sample t-test between urucum and wild-type birds on the red-factor background only.

### Whole genome resequencing, read mapping, and SNP calling

To study the genetic basis of bare-part coloration in canaries, we performed whole-genome resequencing of a DNA pool of urucum individuals (n=20). Briefly, blood was collected into a heparin-free capillary tube and immediately transferred into a vial with 96% ethanol. For DNA sequencing, we extracted genomic DNA from blood using an EasySpin Genomic DNA Kit SP-DT-250 (Citomed), followed by a RNAse A digestion step. DNA quality and purity assessment were performed through spectrophotometry (Nanodrop) and fluorometric quantitation (Qubit dsDNA BR Assay Kit, ThermoScientific). Paired-end sequencing libraries for Illumina sequencing were then generated using the TruSeq DNA PCR-free Library Preparation Kit (Illumina, San Diego, CA) according to the manufacturer protocol, and sequenced using 150 bp paired-end reads on an Illumina instrument (Table S4). Whole genome sequencing data have been deposited in GenBank under the bioproject PRJNA559291.

The reads were trimmed with the program *Trimmomatic* (v. 0.36) (37) using the following parameter settings: TRAILING = 15, SLIDINGWINDOW = 4:20, and MINLEN =30. Sequencing reads were then mapped to the canary reference genome assembly (SCA1; GenBank assembly accession: GCA_000534875.1) (38) with *BWA-MEM* (39), followed by local realignment performed using *GATK RealignerTargetCreator* and *IndelRealigner* options (40). SNP calling was performed using the Bayesian haplotype-based method implemented in *Freebayes* (v0.9.21-26-gbfd9832) (41).

### Genome-wide analysis of genetic differentiation

To identify the genomic region containing the urucum factor, we summarized genetic differentiation between urucum and red lipochrome canaries (data from a previous study (7)) in windows of 20kb moved in steps of 5kb across the genome using the fixation index (*F*_*ST*_). *F*_*ST*_ estimates for each window were obtained by means of an unbiased estimator that corrects for unequal sample sizes across positions due to differences in the depth of coverage, as implemented in the *PoPoolation2* package (42). We utilized the following filters: 1) a per base Phred quality score of 20 or higher; 3) a minimum coverage of 10X per position; 4) a maximum coverage of three times the average coverage per pool; 5) at least four reads supporting the minor allele in polymorphic sites. Only windows that passed our quality filters in ≥30% of positions were kept.

### Crosses and linkage analysis

For linkage analysis, we established crosses between urucum individuals and red lipochrome canaries. The resulting F1 individuals (six pairs) were backcrossed to urucum canaries. From these backcrosses, we obtained 18 individuals and seven expressed red coloration in their beaks and bare-parts (expected to be homozygous for the urucum factor), while 11 were wild type (expected to be heterozygous for the urucum factor). Blood was collected from all individuals and used for DNA extraction as described above. To test for linkage between the candidate genomic region (see Results) and the urucum phenotype in our pedigree, we genotyped a SNP by means of Sanger sequencing of a small amplicon (Table S6). Animal care complied with national and international regulations for the maintenance of live birds in captivity (FELASA, Federation of European Laboratory Animal Science Associations).

### Genotyping for identical-by-descent mapping

We performed identical-by-descent mapping (IBD) by genotyping a panel of 30 SNPs located in the vicinity of our candidate region both in urucum and wild-type individuals. The SNPs were randomly chosen from whole genome resequencing data of several breeds and wild canaries obtained as part of a previous study (GenBank bioproject PRJNA300534 (7)). Blood and DNA was extracted as described above. We genotyped 24 urucum individuals, which are supposed to be homozygous for the causative mutation, and 14 other color canaries belonging to three breeds (white dominant, n=4; lipochrome yellow, n=5; lipochrome red, n=5). Genotyping was carried out using Sequenom (San Diego, USA) iPlex technology and detected on a Sequenom MassArray K2 platform available at the Instituto Gulbenkian de Ciência (Lisbon, Portugal). Sequenom MassARRAY® Assay Design 3.0 software was used to design primers and establish the multiplex conditions. The resulting spectra and plots were manually inspected, and software genotype calls were corrected whenever required. Individual genotypes are available online from the Dryad Digital Repository (XXXXXX).

### SNP/Indel functional annotation

We functionally annotated SNP and indel variants using the genetic variant annotation and effect prediction toolbox *SnpEff* (43). To search for causative mutations within our candidate region, we screened our candidate regions for variants that could potentially alter protein structure and function, such as nonsynonymous, frameshift, STOP, and splice site mutations.

### Detection of structural rearrangements

The identification of structural variants within the candidate region was performed using three approaches: 1) *Breakdancer* (44), which uses read pair orientation and insert size; 2) *DELLY* (45), which uses paired-end information and split-read alignments; and 3) *LUMPY* (46), which uses a combination of multiple signals including paired-end alignment, split-read alignment, and read-depth information. Candidate structural variants were manually inspected using *IGV* (47) and intersected with the canary genome annotation (38).

### Phylogenetic conservation and protein structural modelling

We extracted *BCO2* protein sequences from NCBI and checked for the conservation of the mutated amino acid in other species. We included 175 vertebrate species from a wide of taxonomic groups, including mammals, birds, amphibians, fish and reptiles. Sequences were aligned using *ClustalW* multiple alignment implemented in *BioEdit* v7.0.5 (48).

To infer the impact of the candidate nonsynonymous mutation in the protein structure of *BCO2*, we generated a homology model of canary BCO2 through the Swiss-Model server (49) using bovine RPE65 as the template (PDB accession code: 4RYZ) (50). The R417H substitution and rotamer analysis were performed using *Coot* (51). Images of the wild type and mutant structural models were generated using *Pymol* (Schrödinger, LLC).

### Allelic imbalance

We evaluated whether regulatory variation could explain the urucum phenotype by accessing allelic imbalance in the beak of a F1 individual from a cross between a urucum bird and wild-type individual (i.e. heterozygous for the urucum factor). After dissection, RNA was extracted using the RNeasy Mini Kit (Qiagen), followed by cDNA synthesis using ~1 μg of RNA and the GRS cDNA Kit (GRiSP). To quantify the relative expression of each allele (mutation or wild type), we designed primers to amplify a small fragment from cDNA overlapping the nonsynonymous mutation associated with the urucum phenotype (Table S6). Primers were located on exon-exon junctions to avoid spurious amplification from genomic DNA. Amplification of the cDNA template with 5’ labeled primers was done with the protocol described in (24). Amplicons were sequenced on an Illumina MiSeq system (MiSeq v3 500-cycle kit, 2×250 bp reads). To calculate the relative proportion of alleles expressed in the beak of the heterozygous individual, we mapped the reads with *BWA-MEM* (39) on a reference containing just the *BCO2* cDNA sequence and counted the number of reads corresponding to the reference and alternative alleles.

### Functional analysis of *BCO2* wild-type and mutant proteins

We assayed recombinantly expressed wild-type and mutant *BCO2* enzymes following methods adapted from (52). We amplified the coding sequence of wild-type *BCO2* from the cDNA of a red-factor canary with the primers listed in Table S6 using Phusion Hot Start PCR master mix (M0536, NEB Inc., Ipswich, MA) following the manufacturer’s suggested conditions. We cloned this amplicon into the pET28a vector at the Ned1 and Xho1 restriction sites and confirmed by Sanger sequencing (Eurofins, Louisville, KY). We then introduced the candidate variant into this construct with site-directed mutagenesis by PCR. The primers and procedure for mutagenesis are detailed in Supplementary Materials (Table S6).

We transformed BL21 De3 cells with the wild-type and mutant constructs, grew 1 L liquid cultures of each, and induced protein expression at 16°C overnight (16 hrs) with Isopropyl β-D-1-thiogalactopyranoside following the suggested methods of the vector manufacturer (EMD Biosciences, Madison, WI). We confirmed the expression of the recombinant enzymes by visualizing protein expression in pre- and post-induction cultures on an SDS-PAGE gel. We pelleted bacterial cultures by centrifugation and resuspended pellets in Tricine buffer (50 mM Tricine, 100 mM NaCl, 1% Tween 40) and lysed cells with a cycle of freeze-thaw and brief sonication. We centrifuged the lysed cells at 10,000 g for 30 min at 4°C, collected the supernatant, and measured total protein concentration of the lysates (Pierce™ BCA Protein Assay Kit, ThermoFisher, Waltham, MA). We then used these crude lysates to assayed enzyme activity as described below.

We prepared carotenoid substrates by fractioning all-trans-zeaxanthin and all-trans-canthaxanthin from commercial carotenoid additives (Optisharp, Carophyll Red, DSM Inc, Heerlen, Netherlands) with HPLC. We evaporated the purified carotenoids to dryness under a stream of nitrogen and resuspended by sonication in 2.5 ml of Tricine buffer with 1.6% Tween 40. We combined 400 ul of each carotenoid suspension with a volume of each enzyme lysate containing 1 mg of total protein and supplemented the reaction with 10 μM FeSO4 and 0.3 mM dithiothreitol. We incubated these reactions in the dark at 37°C, shaking at 180 rpm, for 2 hrs. As a negative control, we denatured proteins by boiling the lysates at 100°C for 20 mins and assayed as above. We extracted the reaction products by adding 1 ml of ethanol vortexing briefly, followed by 2 ml of hexane:tert-Butyl methyl ether (1:1), centrifuging briefly and collecting the upper phase. We dried these extracts under a stream of nitrogen and then treated with sodium borohydride to reduce and stabilize apocarotenoid aldehyde products of *BCO2* cleavage. To do this, we resuspended the dried extracts in 1 ml of ethanol, added a few crystals of sodium borohydride, capped reaction tubes with nitrogen, and incubated for 30 mins in the dark. We then added 2 ml of deionized water to the reaction and extracted again with 2 ml of hexane:tert-Butyl methyl ether. We again dried these extracts under a stream of nitrogen and then resuspended in 140 μL of mobile phase (methanol:acetonitrile 1:1). We injected 50 μl of each assay extract into an Agilent 1100 series HPLC fitted with a YMC carotenoid 5.0 μm column (4.6 mm × 250 mm, YMC) cooled to 18°C. We separated pigments with a constant mobile phase flow rate of 1.0◻ml/min and a composition gradient beginning with acetonitrile:methanol (50:50) for 9 min, followed by a ramp up to acetonitrile:methanol:dichloromethane (44:44:12) (vol:vol:vol) from 9–11 min, isocratic through 21 min, then a second ramp up to acetonitrile:methanol:dichloromethane (35:35:30) from 21–26 min followed by isocratic conditions through 35 min. We monitored the sample runs with a photodiode array detector and identified putative cleavage products by comparison to authentic standards (galloxanthin, CaroteNature GmbH, Ostermundigen, Switzerland) or inferred identity from relative retention times and absorbance spectra.

## ACKNOWLEDGEMENTS

This work was supported by the Fundação para a Ciência e Tecnologia (FCT) through POPHQREN funds from the European Social Fund and Portuguese MCTES (FCT Investigator grants to MC [IF/00283/2014/CP1256/CT0012]); and National funds (Transitory Norm contract to RJL [DL57/2016/CP1440/CT0006] by research fellowships attributed to MAG (PD/BD/114042/2015) in the scope of the Biodiversity, Genetics, and Evolution (BIODIV) PhD program at CIBIO/InBIO and University of Porto; and by the project “PTDC/BIAEVL/31569/2017 - NORTE -01-0145-FEDER-30288”, co-funded by NORTE2020 through Portugal 2020 and FEDER Funds, and by National Funds through FCT. MBT received support from the University of Tulsa. PDK was supported by funds from the University of California, Irvine. We also thank of Ibadinis Lda (Versele Laga distributor) for all the support with food supplements and cages. We are grateful for breeder’s help with the samples, namely Alvaro Blasina from Brasil and Susana Mondelo from Spain.

## REFERENCES

1. G. E. Hill, K. J. McGraw, Bird Coloration: voI I, Mechanisms and Measurements. (Harvard University Press, 2006).

2. Mayr E., Animal species and evolution. (Harvard University Press, 1963).

3. M. B. Andersson, Sexual selection (Princeton University Press, 1994).

4. G. Hill, K. J. McGraw, Bird Coloration: Function and Evolution (Vol 2) (Harvard University Press, 2006).

5. J. K. Hubbard, J. A. C. Uy, M. E. Hauber, H. E. Hoekstra, R. J. Safran, Vertebrate pigmentation: from underlying genes to adaptive function. Trends Genet. 26, 231–239 (2010).

6. D. P. L. Toews, N. R. Hofmeister, S. A. Taylor, The Evolution and Genetics of Carotenoid Processing in Animals. Trends Genet. 33, 171–182 (2017).

7. R. J. Lopes, et al., Genetic Basis for Red Coloration in Birds. Curr. Biol., 1427–1434 (2016).

8. N. I. Mundy, et al., Red Carotenoid Coloration in the Zebra Finch Is Controlled by a Cytochrome P450 Gene Cluster. Curr. Biol., 1–6 (2016).

9. M. B. Toomey, et al., High-density lipoprotein receptor *SCARB1* is required for carotenoid coloration in birds. PNAS 114, 5219–5224 (2017).

10. R. E. Koch, K. J. McGraw, G. E. Hill, Effects of diet on plumage coloration and carotenoid deposition in red and yellow domestic canaries (*Serinus canaria*). Wilson J. Ornithol. 128, 328–333 (2016).

11. T. R. Birkhead, K. Schulze-Hagen, R. Kinzelbach, Domestication of the canary, *Serinus canaria* - the change from green to yellow. Arch. Nat. Hist. 31, 50–56 (2004).

12. P. Clement, A. Harris, J. Davis, Finches and sparrows: an identification guide (Princeton University Press, 1993).

13. C. Dela Seña, et al., Substrate Specificity of Purified Recombinant Chicken β-Carotene 9’,10’-Oxygenase (BCO2). J. Biol. Chem. 291, 14609–19 (2016).

14. T. H. Goldsmith, J. S. Collins, S. Licht, The cone oil droplets of avian retinas. Vision Res. 24, 1661–1671 (1984).

15. M. B. Toomey, et al., A complex carotenoid palette tunes avian colour vision. J. R. Soc. Interface 12, 20150563 (2015).

16. M. B. Toomey, J. C. Corbo, Evolution, Development and Function of Vertebrate Cone Oil Droplets. Front. Neural Circuits 11, 97 (2017).

17. M. B. Toomey, et al., Complementary shifts in photoreceptor spectral tuning unlock the full adaptive potential of ultraviolet vision in birds. Elife 5, e15675 (2016).

18. G. B. R. Walker, D. Avon, Coloured, type, and song canaries◻: a complete guide (Blandford, 1993).

19. C. Kiefer, et al., Identification and characterization of a mammalian enzyme catalyzing the asymmetric oxidative cleavage of provitamin A. J. Biol. Chem. 276, 14110–14116 (2001).

20. J. Amengual, et al., A mitochondrial enzyme degrades carotenoids and protects against oxidative stress. FASEB J. 25, 948–959 (2011).

21. J. Eriksson, et al., Identification of the Yellow Skin Gene Reveals a Hybrid Origin of the Domestic Chicken. PLoS Genet. 4, e1000010 (2008).

22. A. Fallahshahroudi, E. Sorato, J. Altimiras, P. Jensen, The Domestic <BCO2> Allele Buffers Low-Carotenoid Diets in Chickens-Possible Fitness Increase Through Species Hybridization. Genetics 212, 1445–1452 (2019).

23. D. P. L. Toews, et al., Plumage Genes and Little Else Distinguish the Genomes of Hybridizing Warblers. Curr. Biol. 26, 2313–2318 (2016).

24. P. Andrade, et al., Regulatory changes in pterin and carotenoid genes underlie balanced color polymorphisms in the wall lizard. Proc. Natl. Acad. Sci., 201820320 (2019).

25. D. P. Kloer, S. Ruch, S. Al-Babili, P. Beyer, G. E. Schulz, The structure of a Retinal-Forming Carotenoid Oxygenase. Science. 308, 267–269 (2005).

26. P. D. Kiser, M. Golczak, D. T. Lodowski, M. R. Chance, K. Palczewski, Crystal structure of native RPE65, the retinoid isomerase of the visual cycle. Proc. Natl. Acad. Sci. 106, 17325–17330 (2009).

27. J. Singh, J. M. Thornton, M. Snarey, S. F. Campbell, The geometries of interacting arginine-carboxyls in proteins. FEBS Lett. 224, 161–171 (1987).

28. N. V. Yanishlieva, K. Aitzetmüller, V. Raneva, β-Carotene and lipid oxidation. Lipid - Fett 100, 444–462 (1998).

29. D. I. Våge, I. A. Boman, A nonsense mutation in the beta-carotene oxygenase 2 (BCO2) gene is tightly associated with accumulation of carotenoids in adipose tissue in sheep (Ovis aries). BMC Genet. 11, 10 (2010).

30. J. Strychalski, P. Brym, U. Czarnik, A. Gugołek, A novel AAT-deletion mutation in the coding sequence of the BCO2 gene in yellow-fat rabbits. J. Appl. Genet. 56, 535–537 (2015).

31. R. Tian, W. S. Pitchford, C. A. Morris, N. G. Cullen, C. D. K. Bottema, Genetic variation in the β, β-carotene-9′, 10′-dioxygenase gene and association with fat colour in bovine adipose tissue and milk. Anim. Genet. 41, 253–259 (2010).

32. E. N. K. Iverson, J. Karubian, The role of bare parts in avian signaling. Auk 134, 587–611 (2017).

33. K. A. Huggins, K. J. Navara, M. T. Mendonça, G. E. Hill, Detrimental effects of carotenoid pigments: the dark side of bright coloration. Naturwissenschaften 97, 637–644 (2010).

34. G. P. Lobo, J. Amengual, G. Palczewski, D. Babino, J. von Lintig, Mammalian Carotenoid-oxygenases: Key players for carotenoid function and homeostasis. Biochim. Biophys. Acta - Mol. Cell Biol. Lipids 1821, 78–87 (2012).

35. G. P. Lobo, A. Isken, S. Hoff, D. Babino, J. von Lintig, *BCDO2* acts as a carotenoid scavenger and gatekeeper for the mitochondrial apoptotic pathway. Development 139(2012).

36. K. J. McGraw, J. Hudon, G. E. Hill, R. S. Parker, A simple and inexpensive chemical test for behavioral ecologists to determine the presence of carotenoid pigments in animal tissues. Behav Ecol Sociobiol 57, 391–397 (2005).

37. A. M. Bolger, M. Lohse, B. Usadel, Trimmomatic: a flexible trimmer for Illumina sequence data. Bioinformatics 30, 2114–2120 (2014).

38. C. Frankl-Vilches, et al., Using the canary genome to decipher the evolution of hormone-sensitive gene regulation in seasonal singing birds. 1–25 (2015).

39. H. Li, R. Durbin, Fast and accurate long-read alignment with Burrows-Wheeler transform. Bioinformatics 26, 589–95 (2010).

40. A. McKenna, et al., The Genome Analysis Toolkit: A MapReduce framework for analyzing next-generation DNA sequencing data. Genome Res. 20, 1297–1303 (2010).

41. E. Garrison, G. Marth, Haplotype-based variant detection from short-read sequencing. arXiv Prepr. arXiv1207.3907, 9 (2012).

42. R. Kofler, R. V. Pandey, C. Schlotterer, PoPoolation2: identifying differentiation between populations using sequencing of pooled DNA samples (Pool-Seq). Bioinformatics 27, 3435–3436 (2011).

43. P. Cingolani, et al., A program for annotating and predicting the effects of single nucleotide polymorphisms, SnpEff. Fly (Austin). 6, 80–92 (2012).

44. K. Chen, et al., BreakDancer: an algorithm for high-resolution mapping of genomic structural variation. Nat. Methods 6, 677–681 (2009).

45. T. Rausch, et al., DELLY: structural variant discovery by integrated paired-end and split-read analysis. Bioinformatics 28, i333–i339 (2012).

46. R. M. Layer, C. Chiang, A. R. Quinlan, I. M. Hall, LUMPY: A probabilistic framework for structural variant discovery. Genome Biol. 15, R84 (2014).

47. J. T. Robinson, et al., Integrative genomics viewer. Nat. Biotechnol. 29, 24–26 (2011).

48. T. A. Hall, BioEdit: a user-friendly biological sequence alignment editor and analysis program for Windows 95/98/NT. Nucleic Acids Symp Ser 41, 95–98 (1999).

49. A. Waterhouse, et al., SWISS-MODEL: homology modelling of protein structures and complexes. Nucleic Acids Res. 46, W296–W303 (2018).

50. J. Zhang, et al., Molecular pharmacodynamics of emixustat in protection against retinal degeneration. J. Clin. Invest. 125, 2781–2794 (2015).

51. P. Emsley, B. Lohkamp, W. G. Scott, K. Cowtan, IUCr, Features and development of *Coot*. Acta Crystallogr. Sect. D Biol. Crystallogr. 66, 486–501 (2010).

52. J. R. Mein, G. G. Dolnikowski, H. Ernst, R. M. Russell, X.-D. Wang, Enzymatic formation of apo-carotenoids from the xanthophyll carotenoids lutein, zeaxanthin and β-cryptoxanthin by ferret carotene-9′,10′-monooxygenase. Arch. Biochem. Biophys. 506, 109–121 (2011).

