## Supplementary materials for "Genetic Basis of *De Novo* Appearance of Carotenoid Ornamentation in Bare-Parts of Canaries"

**Table S1. The concentrations (µg g^-1^ of tissue) of specific carotenoid types in the beak tissues of wild-type (mean ± S.D.) and each urucum canary.**

| **Carotenoid** | **Wild-type red** | **Urucum red #3** | **Urucum red #5** | **Wild-type yellow** | **Urucum yellow #4** |
| --- | --- | --- | --- | --- | --- |
| peak 6.8 | - | 0.16 | 0.08 | 0.01 ± 0.02 | 0.17 |
| peak 7.2 | - | 0.11 | 0.11 | - | 0.03 |
| Lutein | 0.04 ± 0.07 | - | - | 0.08 ± 0.09 | 1.06 |
| *cis*-canthaxanthin | - | 0.36 | 0.77 | - | 0.12 |
| peak 9.5 | - | - | - | - | 0.10 |
| Zeaxanthin | 0.04 ± 0.08 | - | - | - | - |
| Canthaxanthin | 0.14 ± 0.04 | 5.18 | 3.74 | - | 0.08 |
| peak 10.9 | - | - | - | - | - |
| echinenone | - | 0.74 | 0.39 | - | - |
| Total (includes other minor peaks) | 0.22 ± 0.13 | 6.55 | 5.09 | 0.11 ± 0.14 | 1.71 |

**Table S2. The concentrations (µg g^-1^ of tissue) of specific carotenoid types in the feathers of wild-type (mean ± S.D.) and each urucum canary.**

| **Carotenoid** | **Wild-type red** | **Urucum red #3** | **Urucum red #5** | **Wild-type yellow** | **Urucum yellow #4** |
| --- | --- | --- | --- | --- | --- |
| Canary xanthophyll A | 0.55 ± 0.80 | 1.77 | 2.78 | 2.08 ± 1.48 | 1.76 |
| Canary xanthophyll B | 1.65 ± 2.08 | 4.28 | 8.52 | 3.77 ± 2.42 | 4.16 |
| peak 6.8 | 3.39 ± 2.70 | 8.88 | 13.40 | 0.63 ± 0.19 | 0.29 |
| peak 7.2 | 1.45 ± 0.38 | 2.69 | 2.70 | 0.30 ± 0.24 | 0.00 |
| Lutein | 2.82 ± 1.07 | 5.47 | 4.34 | 0.26 ± 0.05 | 0.50 |
| *cis*-canthaxanthin | 15.26 ± 5.80 | 28.97 | 14.33 | - | - |
| canthaxanthin | 72.98 ± 22.33 | 123.87 | 69.99 | - | - |
| echinenone | 10.68 ± 4.03 | 13.20 | 10.45 | - | - |
| Total (includes other minor peaks) | 111.34 ± 33.15 | 189.13 | 126.51 | 7.61 ± 4.71 | 7.17 |

**Table S3. The concentrations (µg g^-1^ of tissue) of specific carotenoid types in the retinas of wild-type (mean ± S.D.) and each urucum canary.**

| **Carotenoid** | **Wild-type red** | **Urucum red #3** | **Urucum red #5** | **Wild-type yellow** | **Urucum yellow #4** |
| --- | --- | --- | --- | --- | --- |
| *cis*-astaxanthin | 20.40 ± 3.65 | 4.50 | 4.40 | 22.14 ± 4.95 | 37.10 |
| astaxanthin | 141.20 ± 76.17 | 5.40 | 26.50 | 173.29 ± 65.97 | 172.20 |
| peak 8.6 | 23.20 ± 7.12 | 72.60 | 5.00 | 15.43 ± 7.48 | 4.20 |
| peak 9.2 | 32.80 ± 12.07 | 5.50 | 76.20 | 35.57 ± 9.43 | 33.50 |
| canthaxanthin | 22.40 ± 6.80 | 266.30 | 504.20 | - | - |
| peak 4.8 | 18.60 ± 13.76 | 1.00 | - | - | - |
| dihydrogalloxanthin | 64.40 ± 24.54 | 11.00 | - | 150.86 ± 53.82 | - |
| peak 6.5 | 10.00 ± 4.53 | 2.00 | - | 26.29 ± 9.74 | - |
| peak 9.3 | - |  |  | - | - |
| 3’-epilutein | 2.00 ± 2.83 | - | - | 20.00 ± 7.96 | 30.10 |
| Lutein | 9.20 ± 3.11 | - |  | 44.57 ± 22.01 | 14.10 |
| Zeaxanthin | 9.60 ± 6.84 | - | - | 33.29 ± 14.10 | 56.00 |
| peak 25.8 | - | - | - | - | 5.40 |
| peak 27.2 | - | - | - | - | 10.20 |
| peak 27.6 | - | - | - | - | 26.20 |
| Total (includes other minor peaks) | 460.00 ± 140.18 | 420.60 | 628.90 | 647.14 ± 222.39 | 460.00 |

**Table S4. Whole genome resequencing details and read mapping statistics for the two pools used for *F_ST_* analysis.**

| **Breed/Population** | **Bare-part coloration** | **Number of individuals pooled** | **Number of reads^a^** | **Percentage of mapped reads^b^** | **Percentage of reads MQ >=20**^c^ | **Percentage of positions >=1 reads^d^** | **Coverage^e^** |
| --- | --- | --- | --- | --- | --- | --- | --- |
| Lipochrome red | No | 16 | 202,038,581 | 99.24 (92.31) | 88.42 | 96.02 | 16.9X |
| Urucum | Yes | 20 | 222,250,123 | 99.31 (92.64) | 85.54 | 99.97 | 26.5X |

^a^After trimming using *Trimmomatic*.

^b^Percentage of properly paired reads is given in parentheses.

^c^Reads with a Phred score mapping quality (MQ) equal to or greater than 20.

^d^Percentage of positions with at least one mapped read.

^e^Including positions with zero mapped reads.

**Table S5. List of protein-coding genes located with the candidate region for urucum phenotype (*located in IBD region).**

| **Gene** | **Location** | **Orientation** | **Description** |  |
| --- | --- | --- | --- | --- |
| *MPZL2* | NW_007931177:500997-504867 | - | Myelin Protein Zero Like 2 |  |
| *CD3E* | NW_007931177:507652-511777 | + | CD3e Molecule |  |
| *LOC103822979* | NW_007931177:512949-514539 | - | T-cell surface glycoprotein CD3 gamma chain-like |  |
| *ZW10* | NW_007931177:515096-522436 | + | Zw10 Kinetochore Protein |  |
| *DRD2* | NW_007931177:569855-576921 | + | Dopamine Receptor D2 |  |
| *ANKK1* | NW_007931177:590507-602622 | - | Ankyrin Repeat And Kinase Domain Containing 1 |  |
| *TTC12* | NW_007931177:610371-625839 | - | Tetratricopeptide Repeat Domain 12 |  |
| *NCAM1* | NW_007931177:631156-819387 | - | Neural Cell Adhesion Molecule 1 | * |
| *PTS* | NW_007931177:831089-834436 | - | 6-Pyruvoyltetrahydropterin Synthase | * |
| *BCO2* | NW_007931177:836313-850746 | - | Beta-Carotene Oxygenase 2 | * |
| *TEX12* | NW_007931177:855386-856620 | - | Testis Expressed 12 | * |
| *IL18* | NW_007931177:861542-864199 | + | Interleukin 18 | * |
| *SDHD* | NW_007931177:865343-868732 | - | Succinate Dehydrogenase Complex Subunit D | * |
| *LOC103822986* | NW_007931177:870092-872020 | - | NKAPD1 NKAP domain containing 1 | * |
| *DLAT* | NW_007931177:874163-882973 | - | Dihydrolipoamide S-Acetyltransferase | * |
| *DIXDC1* | NW_007931177:886262-908100 | - | DIX Domain Containing 1 |  |
| *LOC103822987* | NW_007931177:913599-918479 | - | CUNH11orf52 chromosome unknown C11orf52 homolog |  |
| *HSPB2* | NW_007931177:922487-924072 | - | Heat Shock Protein Family B (Small) Member 2 |  |
| *CRYAB* | NW_007931177:926301-928409 | + | Crystallin Alpha B |  |
| *FDXACB1* | NW_007931177:931323-934952 | + | Ferredoxin-Fold Anticodon Binding Domain Containing 1 |  |
| *ALG9* | NW_007931177:935845-953645 | + | ALG9 Alpha-1,2-Mannosyltransferase |  |
| *SIK2* | NW_007931177:968384-995914 | - | Salt Inducible Kinase 2 |  |
| *LAYN* | NW_007931177:1000111-1008095 | - | Layilin |  |
| *BTG4* | NW_007931177:1017880-1021291 | + | BTG Anti-Proliferation Factor 4 |  |
| *POU2AF1* | NW_007931177:1078714-1089769 | + | POU Class 2 Homeobox Associating Factor 1 |  |
| *NFKBID* | NW_007931177:1092851-1102554 | + | NFKB Inhibitor Delta |  |
| *COLCA2* | NW_007931177:1107643-1114209 | - | Colorectal Cancer Associated 2 |  |
| *LOC103823036* | NW_007931177:1117735-1125332 | - | similar to RGD1562914 (predicted) |  |
| *HINFP* | NW_007931177:1126635-1131935 | + | Histone H4 Transcription Factor |  |
| *ABCG4* | NW_007931177:1136837-1145092 | + | ATP Binding Cassette Subfamily G Member 4 |  |
| *NLRX1* | NW_007931177:1150756-1155964 | + | NLR Family Member X1 |  |
| *PDZD3* | NW_007931177:1157287-1160314 | + | PDZ Domain Containing 3 |  |
| *CBL* | NW_007931177:1162741-1196504 | + | Cbl Proto-Oncogene |  |
| *MCAM* | NW_007931177:1201344-1218819 | - | Melanoma Cell Adhesion Molecule |  |
| *RNF26* | NW_007931177:1219052-1220085 | + | Ring Finger Protein 26 |  |
| *C1QTNF5* | NW_007931177:1224879-1227445 | - | C1q And TNF Related 5 |  |
| *MFRP* | NW_007931177:1230124-1233732 | - | Membrane Frizzled-Related Protein |  |
| *USP2* | NW_007931177:1235956-1246094 | - | Ubiquitin Specific Peptidase 2 |  |
| *THY1* | NW_007931177:1253201-1256644 | - | Thy-1 Cell Surface Antigen |  |
| *PVRL1* | NW_007931177:1374817-1417137 | - | Nectin Cell Adhesion Molecule 1 |  |

**Table S4. Detailed information about all primers used in this study*.***

|  | **Primer 1** | **Primer 2** | **Extension primer**  **(only for genotyping)** |
| --- | --- | --- | --- |
| **Genotyping** |  |  |  |
| RB_177770364 | ACGTTGGATGTGTGTGTCCCTGAGAACAGC | ACGTTGGATGCTCAGATAAGGCAAGGGCAG | CCACATCCCACACAA |
| RB_177803809 | ACGTTGGATGTTTGAAGTCCTGTCCACAAG | ACGTTGGATGGGTTTTTCACCAAAGCCCTC | CCAGTGGAACAGACC |
| RB_177866964 | ACGTTGGATGCTCTGTTCACACAACACTGG | ACGTTGGATGTTAAGAGCACTTCCTTGCCC | cGCCCCCAGAAGTGTA |
| RB_1771138095 | ACGTTGGATGAAAGGTGAAATGACGAGCCG | ACGTTGGATGGGAGAGCCTTAACAGAACAG | cACAGAACAGGGAGGG |
| RB_177849620 | ACGTTGGATGGAGTCCAGAGGGCACAAATC | ACGTTGGATGACAACATGGAAGGCAAGCAG | CCCGACCCCAGCCCAAA |
| RB_177642749 | ACGTTGGATGAGGAAGGGCTGTGTCCTTGT | ACGTTGGATGGGCCAAAATGACAAAACCCC | AAAGGCAGCAATAACCA |
| RB_177830073 | ACGTTGGATGAATGAGGGCCATTCCCTTAG | ACGTTGGATGCACCCCCTTTGACAAGGAAA | gGGAAAAGGATCCCCCA |
| RB_177927503 | ACGTTGGATGTGTGAACCTGGATGTGAAGC | ACGTTGGATGTTCCCGTGGATCTCGATCAT | aacCTCGATCATGTCCCC |
| RB_177821659 | ACGTTGGATGTCTGTCTTTTCTTTGCTCCC | ACGTTGGATGAATCCTCTTTCTCTCCCCAG | TATCTTTTTGCCCGAACT |
| RB_177555773 | ACGTTGGATGGAGAACTTGGAGCTATCACC | ACGTTGGATGTTTGCTGGAGCTCCAGATTG | CTAGGAGGGAGTTTAGAC |
| RB_177874443 | ACGTTGGATGAATGACAGAGTGACCAGGAG | ACGTTGGATGACAGAATAACAGGATCAGCC | GGATCAGCCTGAACAACTT |
| RB_177793924 | ACGTTGGATGACATCTGTCATCTCCAGCAC | ACGTTGGATGAATAGCTCTGAAGTGGCTGG | AACCTCGCATTTCCTATCAC |
| RB_177941117 | ACGTTGGATGTCAAGGATGATGTTCTGGGC | ACGTTGGATGGTGAACTGAGACAGAACTGC | ctgcCCCCACTGCAGGAGAG |
| RB_1771328744 | ACGTTGGATGCTAAAAGGAGCCAGTCCTTC | ACGTTGGATGTTTCCTCCCTTTAAGCACCC | gaaaCCAGAGCTGCAGAAAA |
| RB_1771054485 | ACGTTGGATGGGGATTTCTCACCTTGCCTG | ACGTTGGATGAGATGCCAGTGCGGGAATTG | aGGGAATTGCTGAGGAAAAT |
| RB_1771449572 | ACGTTGGATGGTCAGCAGGGATTTTTCAGG | ACGTTGGATGCTCCTTTTTTGTGAGCAGCC | gtgtAGCCGCAGCCTCAGCCA |
| RB_177889716 | ACGTTGGATGAGTGTCCAACACTTGTGCAG | ACGTTGGATGTGTCCCAGAGCACACAGGTT | ggaaCAGGGACCAGAGGACAA |
| RB_1771002052 | ACGTTGGATGTTGCCACAGCTGGAAAGAAG | ACGTTGGATGCCTCTCAGCCTCACTTTTTG | TCAGCCTCACTTTTTGATTATT |
| RB_1771228752 | ACGTTGGATGGTGTTCTGCCATCCCCTTG | ACGTTGGATGCTATAGCAGAAAGTGGGAGC | ggtTGCCCCGACTGACAGGAGA |
| RB_177836252 | ACGTTGGATGTGGGCATCCTTTGTGAACTG | ACGTTGGATGCCACTGCTTGAGCTACAGTT | GCTTGAGCTACAGTTTTTTTACT |
| **Genotyping missense mutation in BCO2** | |  |  |
| GenotPedigree | CTGTGAGAGCAGTGCTGACC | GAAATGTGCAGGGTTGGTTC | NA |
| **Allelic imbalance in BCO2** | |  |  |
| ABalance | CGCCTTCCACCAGATCAAC | TCTTGGTGTCCACATCAACCT | NA |
| **Cloning and mutagenesis of *BCO2*** | |  |  |
| SeCa_BCO2 | CGTACATatgttcgccaaaatcc | CGTACTCGAGgtgggcagtgaagatgc |  |
| Seca_mut-PmII | Ttccacgtgaacaagca | NA | NA |
| SeCa_BCO2_SD_antisense | cgtccagagggagaacaaagtggcgagggaaagctc | NA | NA |
| SeCa_BCO2_SD_sense: | Gagctttccctcgccactttgttctccctctggacg | NA | NA |

**Figure S1. Representative HPLC chromatograms of carotenoid extracts from the beak tissue wild-type and urucum canaries. The UV/Vis spectra of each labeled peak are presented in figure S3.**

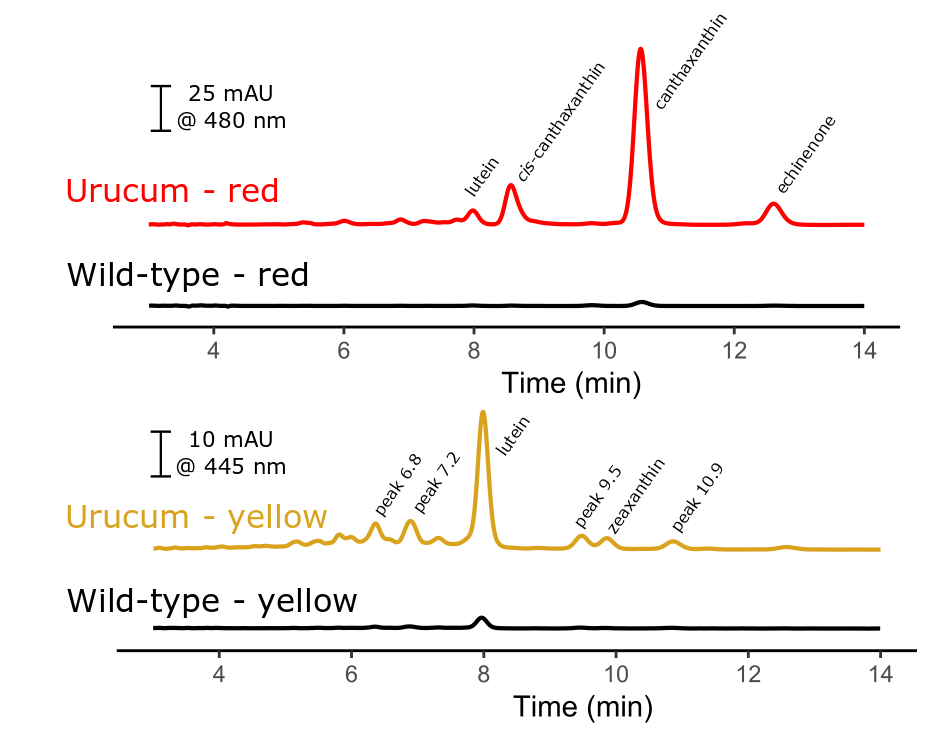

**Figure S2. Representative UV/Vis light absorbance spectra, normalized to absorbance maximum, of the major carotenoids observed in extracts from the beak tissues and feathers of wild-type and urucum canaries.**

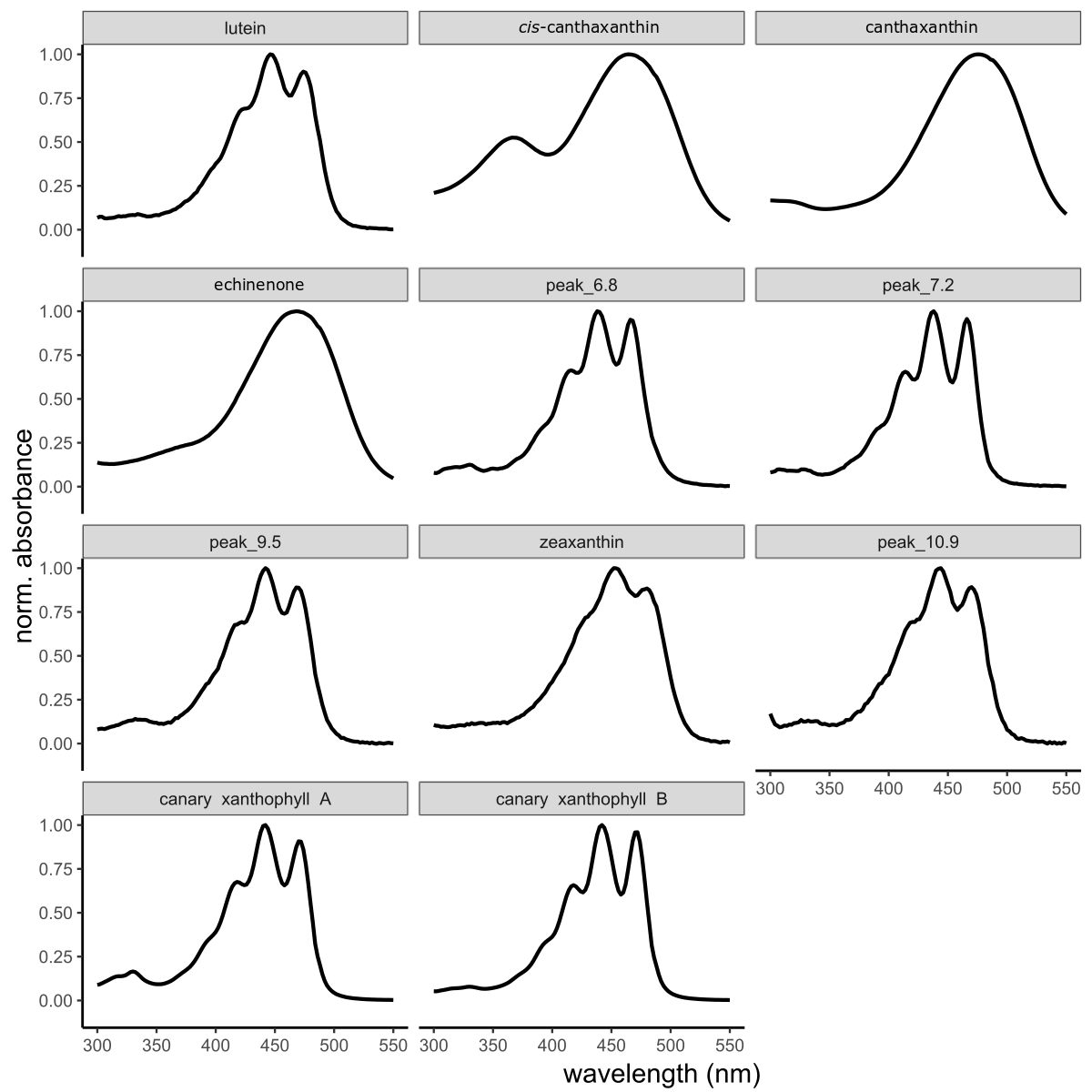

**Figure S3. Representative HPLC chromatograms of carotenoid extracts from the feathers of wild-type and urucum canaries. The UV/Vis spectra of each labeled peak are presented in figure S2.**

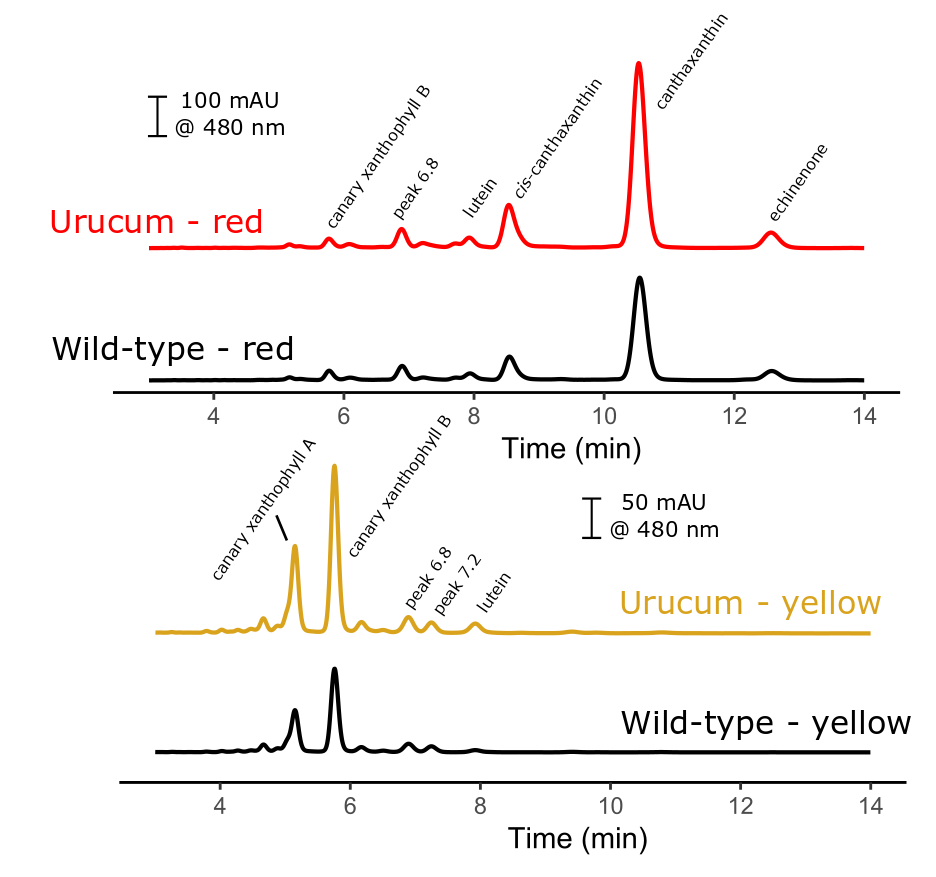

**Figure S4. Representative HPLC chromatograms of retinal carotenoid extracts from wild-type and urucum canaries. The (A) ketocarotenoid, (B) other xanthophyll, and (C) apocarotenoid components were measured at differ wavelength and are presented separately. The UV/Vis spectra of each labeled peak are presented in Figure S5.**

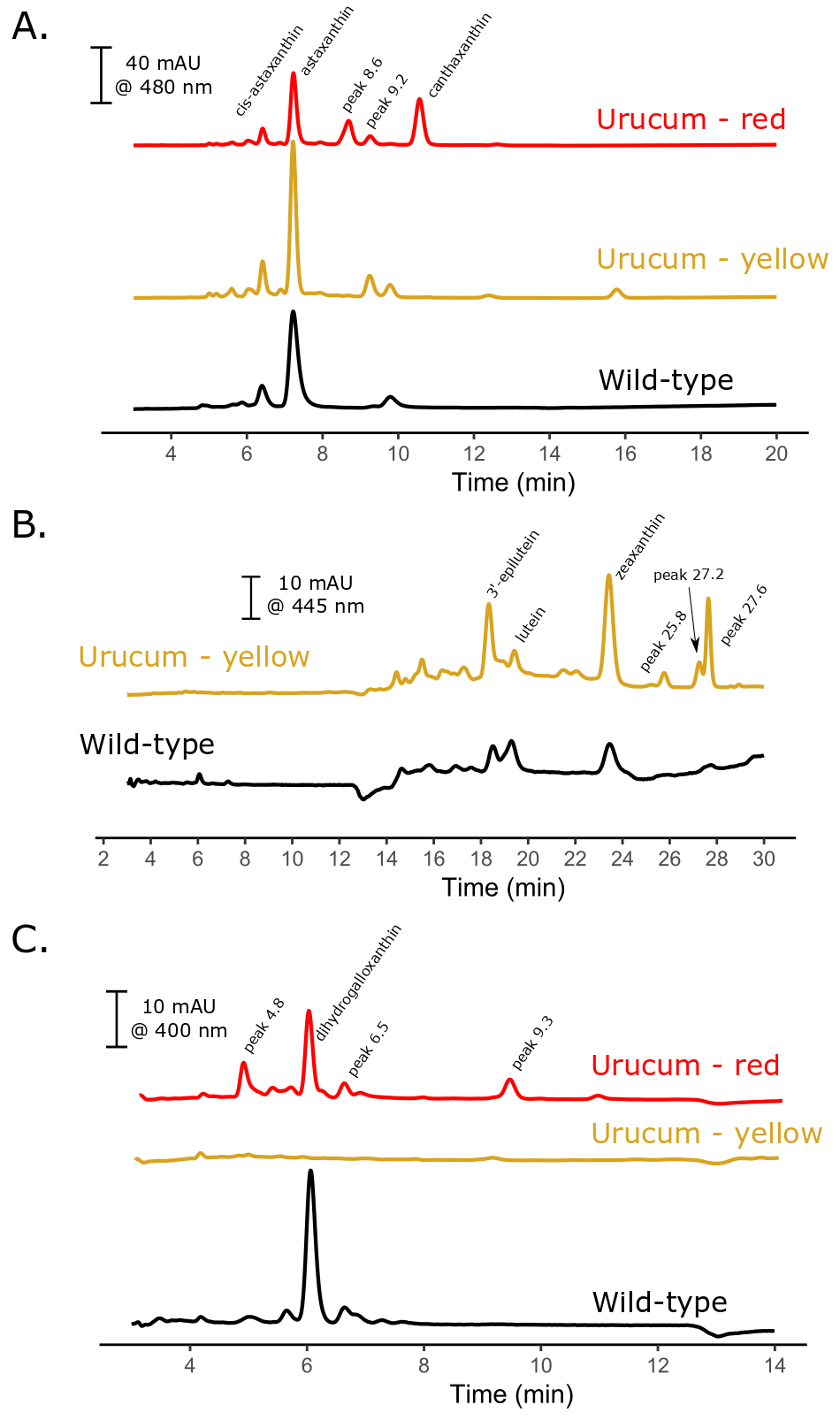

**Figure S5. Representative UV/Vis light absorbance spectra, normalized to absorbance maximum, of the major (A) ketocarotenoid, (B) other xanthophyll, and (C) apocarotenoid components of the retinas of wild-type and urucum canaries.**

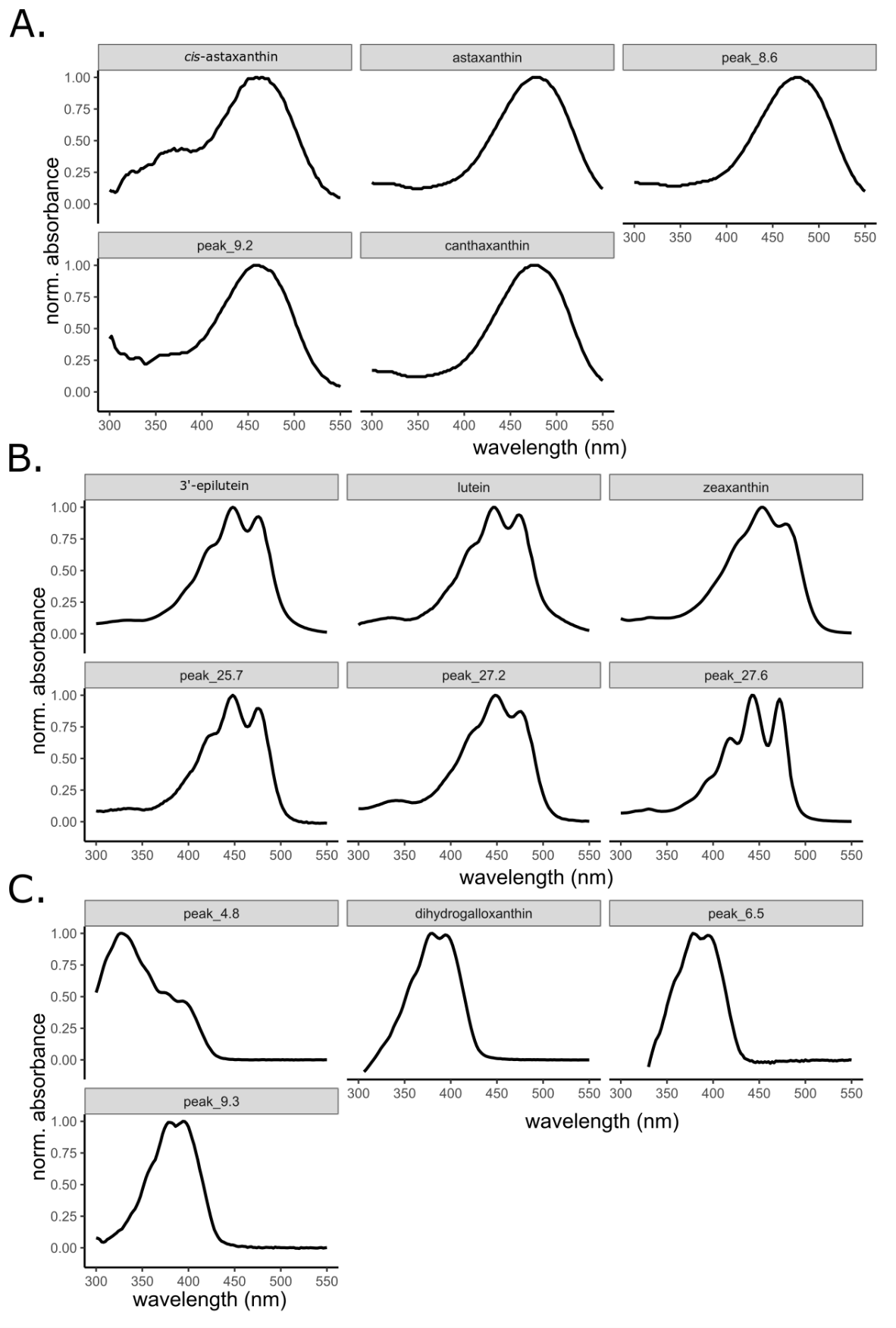

**Figure S6. Retinal canthaxanthin accumulation patterns among urucum and wild-type canaries. Points indicate the value for each individual sampled. The concentration of canthaxanthin is given per gram of total protein in sample.**

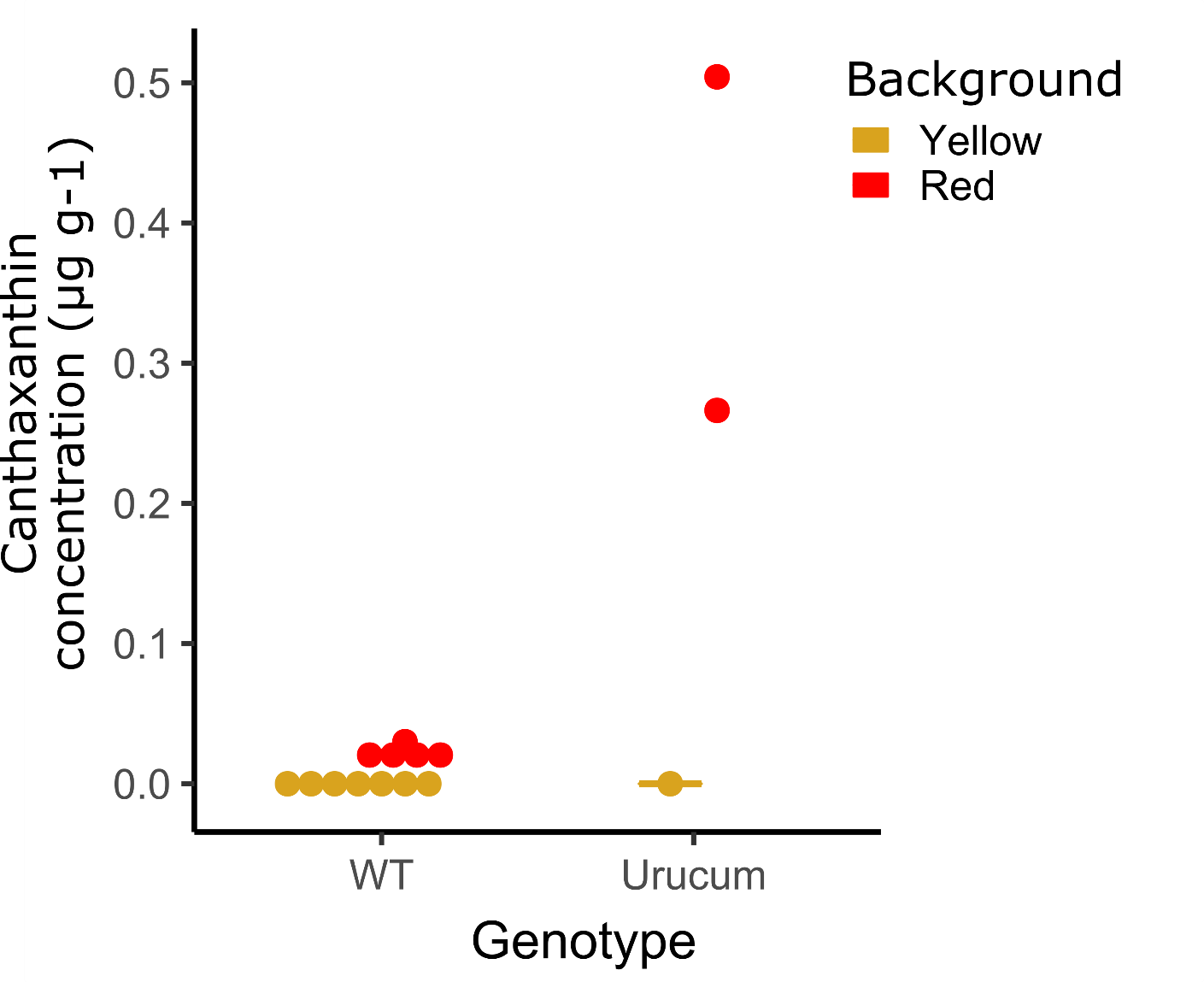

**Figure S7. Urucum canaries have a mutation in the highly conserved exon 9 of *BCO2* (the missense mutation is highlighted in red). Multi-species alignment of vertebrates surrounding the candidate missense mutation in *BCO2*. The dots represent the same nucleotide as the reference wild-type canary allele at a given position.**
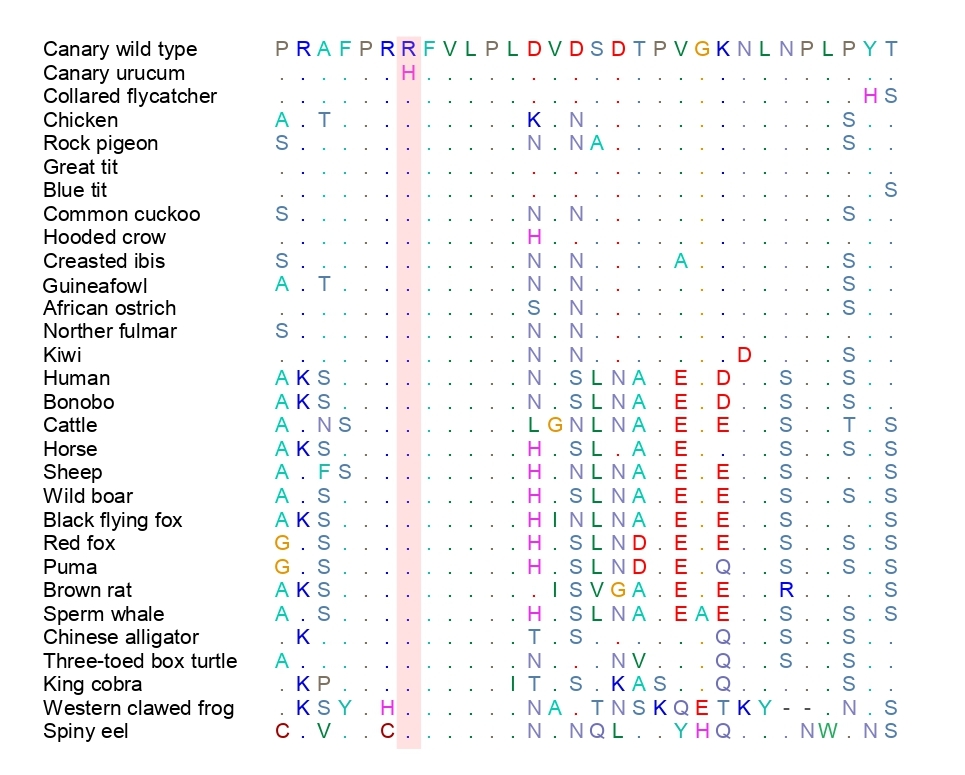

**Figure S8. (A) Representative HPLC chromatograms of carotenoids extracted from reactions containing zeaxanthin substrates and wild-type, mutant, or wild-type heat denatured bacterial lysates. (B) UV-Vis light absorbance spectra of peaks 1 and 2. The short retention time and short-wavelength shifted absorbance spectrum of peak 1 matches an authentic standard of the apocarotenoid galloxanthin (10’-Apo-β-carotene-4,10’-ol). Trace amounts of galloxanthin are also present in the mutant or denatured *BCO2* assays but are likely the product of non-enzymatic oxidation of the zeaxanthin substrate.**

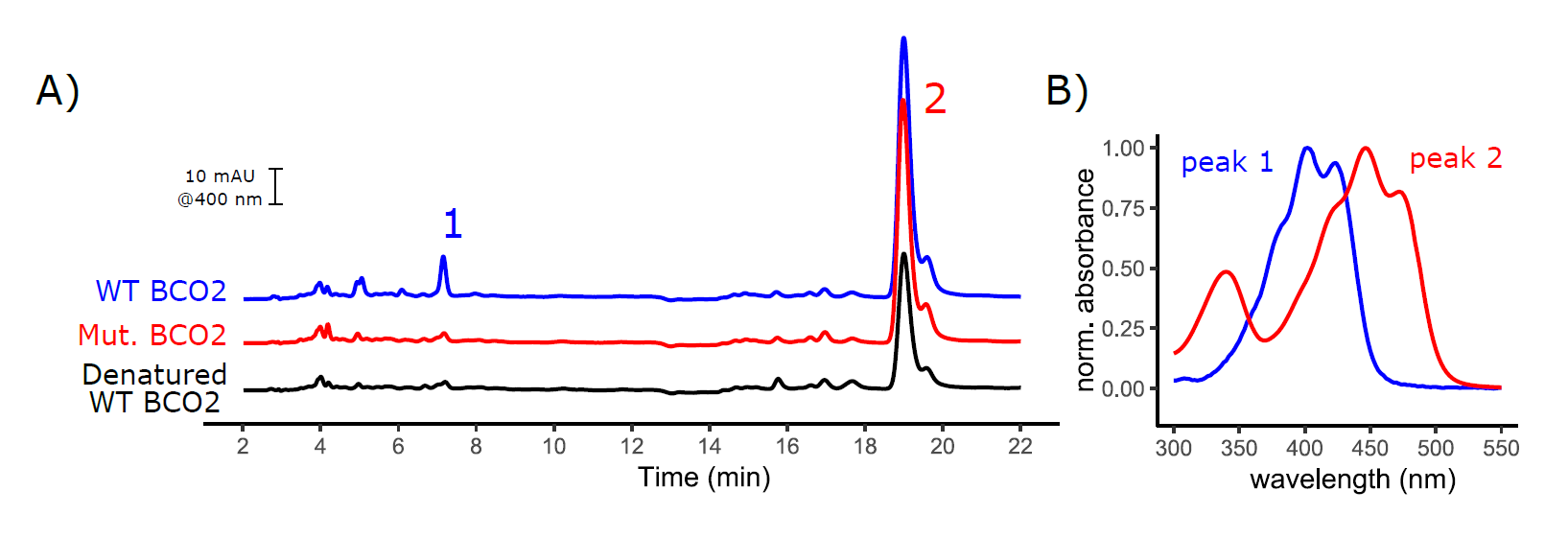
